# Generating high quality assemblies for genomic analysis of transposable elements

**DOI:** 10.1101/2020.03.27.011312

**Authors:** Filip Wierzbicki, Florian Schwarz, Odontsetseg Cannalonga, Robert Kofler

## Abstract

The advent of long-read sequencing holds great promise for research on transposable elements (TEs). Long reads may finally allow us to obtain reliable assemblies of repetitive regions, and thus shed light on many open questions in TE biology, such as the evolution of piRNA clusters, i.e., the master loci controlling TE activity. Currently, many different assembly strategies exist and it is not clear how to obtain the most suitable assemblies for TE research. In fact, it is not even clear how to best identify suitable assemblies as classic quality metrics such as BUSCO and NG50 are ignorant of TEs. To address these problems, we introduce four novel quality metrics that assess i) how well piRNA clusters are assembled (CUSCO) and ii) to which extent an assembly captures the TE landscape of an organism (TE abundance, SNPs and internal deletions). Using these novel metrics, we evaluate the effect of assemblers, polishing, read length, coverage, residual polymorphisms, and finally, identify suitable assembly strategies. Using an optimized approach, we provide high-quality assemblies for the two *Drosophila melanogaster* strains Canton-S and Pi2. Around 80% of the piRNA clusters were contiguously assembled in these two strains. Such high-quality assemblies will provide novel insights into the biology of TEs. It is, for example, an open question of whether piRNA clusters contain abundant presence/absence polymorphism of TE insertions, as expected when piRNA clusters are responsible for stopping TE invasions. A comparison of the sequences of our assembled piRNA clusters reveals that such polymorphisms are indeed abundantly found in clusters.

## Introduction

Transposable elements (TEs) are short stretches of DNA that multiply within genomes. The genomes of most organisms, ranging from mammals to bacteria, are riddled with transposable elements (TEs) (Biémont and Vieira, 2006; Wicker et al., 2007). For example, a striking 80% of the maize genome consists of TEs (Schnable et al., 2009). As TEs are mostly deleterious to hosts (Nuzhdin, 1999), their widespread distribution was long thought to be a mystery. The issue was resolved by two classical works, which suggested that TEs are genomic parasites (Orgel and Crick, 1980; Doolittle and Sapienza, 1980). By multiplying in genomes, TEs enhance their transmission, even if this reduces the fitness of the hosts (Hickey, 1982). However, an unconstrained proliferation of TEs could lead to an accumulation of deleterious TE insertions that may eventually drive host populations to extinction (Brookfield and Badge, 1997). To counteract the selfish activity of TEs, host species evolved sophisticated defense mechanisms that frequently involve small RNAs (Blumenstiel, 2011; Marí-Ordóñez et al., 2013; Goodier, 2016; Czech and Hannon, 2016; Kofler et al., 2018). In mammals and invertebrates, the defense against TEs is based on piRNAs, i.e., small RNAs ranging in size from 23-29nt (Aravin et al., 2007; Gunawardane et al., 2007; Brennecke et al., 2007; Ernst et al., 2017). These piRNAs bind to PIWI-clade proteins and repress TEs at the transcriptional and post-transcriptional level (Gunawardane et al., 2007; Brennecke et al., 2007; Sienski et al., 2012; Le Thomas et al., 2013). piRNAs are derived from discrete genomic loci, the piRNA clusters (Brennecke et al., 2007). These piRNA clusters are likely of central importance for controlling the spread of TEs since it is assumed that a newly invading TE is stopped when one copy of the TE jumps into a piRNA cluster, which triggers the production of piRNAs against the TE that silence it (Bergman et al., 2006; Malone and Hannon, 2009; Zanni et al., 2013; Goriaux et al., 2014; Yamanaka et al., 2014; Ozata et al., 2019; Duc et al., 2019). This view is known as the trap model since piRNA clusters act as genomic traps for active TEs (Bergman et al., 2006).

Apart from causing deleterious effects to hosts, TEs are major drivers of genome evolution (Kazazian, 2004) and may contribute to the adaptation of organisms to novel environments (Daborn et al., 2002; González et al., 2008; Casacuberta and González, 2013). Despite the importance of TEs, many questions about TE biology remain open. This is mainly because most currently available information about TEs is based on short reads, which are too short for resolving repetitive regions unambiguously. Therefore, short reads solely provide an incomplete picture of the TE landscape. The advent of long-read sequencing holds great promise for TE research as long reads will finally enable generating accurate assemblies of repetitive regions. Such high-quality assemblies will enable us to address many important questions in TE biology such as i) what is the genomic distribution and population frequency of different TE variants? ii) are full-length insertions more deleterious than internally deleted ones (full-length TEs could have a lower population frequency than internally deleted ones) (Petrov et al., 2003, 2011)? iii) do similar TE variants cluster on the same chromosomes as expected when TEs proliferate locally (Delattre et al., 2000)? and iv) can we infer the history of TE activity of any organism by investigating networks of nested TE insertions (Bergman et al., 2006)? Most importantly, however, high-quality assemblies will allow us to study the evolution of piRNA clusters. We could, for example, test the hypothesis that the composition of piRNA clusters evolves rapidly, as expected under the widely held trap model (Kelleher et al., 2018; Kofler, 2019). Long-read based assemblies may thus revolutionize TE research. Unfortunately, it is not clear which assembly strategies yield the best assemblies, since many different assembly tools, polishing strategies, sequencing data, and scaffolding approaches could be used. In fact, it is not even clear on how to best identify the most suitable assemblies, as classic quality metrics such as BUSCO and NG50 are ignorant of TEs: BUSCO (Benchmarking Universal Single-Copy Orthologs) provides the fraction of correctly assembled core genes (Simão et al., 2015; Waterhouse et al., 2017) and NG50 gives the size of the smallest contig out of the largest contigs that account for 50% of the reference genome (Earl et al., 2011).

Here, we address these challenges by first introducing four novel quality metrics that assess the quality of an assembly from the perspective of TE biology. These novel quality metrics were used to evaluate the effect of different assembly algorithms, polishing approaches, read lengths, coverages, residual polymorphisms, and scaffolding approaches. Based on these results, we identify strategies that generate high-quality assemblies for TE research. Using such an optimized strategy, we provide novel assemblies for the *Drosophila melanogaster* strains Canton-S and Pi2. Investigating the composition of piRNA clusters in Pi2 and Canton-S, we found that piRNA clusters carry abundant presence/absence polymorphism of TE insertions as predicted by the trap model (Kofler et al., 2018; Kofler, 2019).

## Results

### Assembly quality from the perspective of TE biology

Here, we aim to identify strategies that allow generating assemblies of high quality, suitable for genomic analyses of TEs. In particular, assemblies should i) provide contiguous sequences of piRNA clusters and ii) accurately reproduce the TE landscape, i.e., the abundance and diversity of TEs. We evaluated different assembly strategies with *D. melanogaster*, an organism widely used for studying TE biology (Brennecke et al., 2007; Barrón et al., 2014; Kofler et al., 2015; Moon et al., 2018). We first sequenced the *D. melanogaster* strain Canton-S with i) the Oxford Nanopore long-read technology (coverage 150x, mean read length *≈* 7 kb), ii) Illumina paired-end sequencing (coverage 30x, read length 125 bp) and iii) Hi-C (coverage 530x, read length 125 bp) (supplementary table 1). Since commonly used assembly quality metrics, such as BUSCO and NG50 (Earl et al., 2011; Simão et al., 2015), are ignorant of TEs, we developed four novel quality metrics. With these metrics, we aim to estimate the contiguity of piRNA clusters and the accuracy of the assembled TE landscape.

We first developed a novel metric that allows us to assess whether piRNA clusters, the master regulators of TE activity, are contiguously assembled. In essence, the CUSCO value (*C* luster B*USCO*) estimates the fraction of contiguously assembled piRNA clusters (fig. 1A). Based on the reference genome of *D. melanogaster*, we identified flanking sequences for 85 out of the 142 annotated piRNA clusters of *D. melanogaster* (Brennecke et al., 2007) (available as supplementary data). Flanking sequences close to piRNA clusters were preferred. Note that we excluded clusters that were annotated at the ends of chromosomes (10) and on the highly fragmented U-chromosome (46), as flanking sequences can not be obtained for these clusters. These flanking sequences may then be mapped to an assembly of interest. We assume that a piRNA cluster is contiguously assembled when both flanking sequences align to the same contig/scaffold. The CUSCO value is the fraction of flanking sequences aligning to the same contig/scaffold, and thus a proxy for the fraction of contiguously assembled piRNA clusters (fig. 1A). Depending on whether poly-N sequences (i.e., gaps in the assembly) are tolerated between the flanking sequences, a contig-CUSCO (without gaps), or a scaffold-CUSCO (tolerating gaps) can be computed (fig. 1A). We ignored the length of piRNA clusters for computing CUSCO values as theoretical work suggests that piRNA clusters could be highly polymorphic: abundant presence/absence polymorphism of TE insertions in piRNA clusters may render the length of the clusters highly variable among individuals (Kelleher et al., 2018; Kofler, 2019). To illustrate the usage of the CUSCO, we generated two assemblies of Canton-S: i) an assembly based on Illumina short reads (ABySS (Simpson et al., 2009)) and ii) an assembly based on ONT long reads (Canu (Koren et al., 2017))) and several rounds of polishing using the long and the short reads (3x Racon and 3x Pilon (Walker et al., 2014; Vaser et al., 2017)). For both short and long reads, the coverage was 30x. CUSCO values differed substantially between the short- and long-read based assemblies (fig. 1B). A mere 5.88% of piRNA clusters were contiguously assembled with the short reads, while 60% were contiguously assembled with long reads. As we did not perform scaffolding, only the contig-CUSCO was calculated (fig. 1B). By contrast, both assemblies show high BUSCO values, which illustrates that BUSCO is of limited use for estimating the suitability of assemblies for genomic analyses of TEs (fig. 1B).

**Figure 1:**
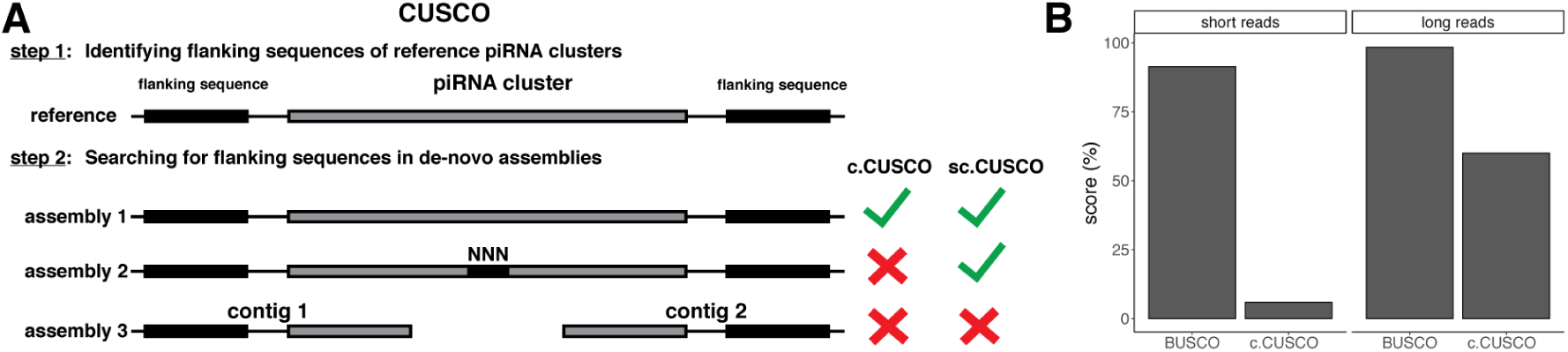
The CUSCO value (<u>C</u>luster B<u>USCO</u>) estimates the fraction of correctly assembled piRNA clusters. A) For computing CUSCO values, it is necessary to identify unique sequences flanking piRNA clusters. By aligning the flanking sequences back to an assembly of interest, the CUSCO value can be computed as the percentage of contiguously assembled piRNA clusters. Depending on whether or not poly-N sequences, i.e., assembly gaps, are tolerated between the flanking sequences, a contig-CUSCO (c.CUSCO) and a scaffold-CUSCO (sc.CUSCO) may be distinguished. B) BUSCO and CUSCO values for different assemblies of Canton-S (30x coverage for short and long reads). Although long- and short-read based assemblies have similar BUSCO values, CUSCO values differ substantially.

To estimate whether an assembly accurately reproduces the TE landscape, we measured, for each TE family, three different features: 1) the abundance (in reads per million: rpm), 2) the number of single nucleotide polymorphisms (SNPs), and 3) the number of internal deletions (IDs). A comparison of the expected and observed values for these three features will allow us to estimate the quality of an assembly. The key idea is that the expected TE landscape can be directly derived from the Illumina raw reads using our novel tool, DeviaTE (Weilguny and Kofler, 2019). Briefly, DeviaTE aligns Illumina reads to the consensus sequences of TEs and provides estimates of the abundance (rpm) and diversity (SNPs and IDs counts) of each TE family (for example, see supplementary fig. 1). The observed TE landscape can be computed for each assembly by using DeviaTE with artificial reads derived from the assembly. To avoid biases and sampling noise, these artificial reads should be uniformly distributed across the assembly and have the same length as the Illumina raw reads used for inferring the expected TE landscape. This approach yields the expected and the observed abundance and diversity of each TE family (for example, see supplementary fig. 2). To summarize these results across all TE families (e.g., 127 TE families in *D. melanogaster*), we perform a linear regression between the expected and the observed values (fig. 2). We propose to use the slope of the regression line as a novel quality metric. This yields, in total, three novel quality metrics (slope of abundance, SNP count, and ID count) that estimate how well an assembly captures the TE landscape. High-quality assemblies that accurately reproduce the TE landscape will have regression slopes of 1.0 for each of the three features. Assemblies that, for example, overestimate the TE abundance, will have a slope *>* 1.0 and assemblies that underestimate the TE abundance a slope *<* 1.0. We again illustrate the usage of these metrics with the short- and the long-read based assemblies of Canton-S. The short-read based assembly poorly reproduced the TE landscape (fig 2). The abundance of many families was underestimated, and the diversity of many TE families (SNPs and IDs) was overestimated (fig. 2). By contrast, the long-read based assembly captured the TE landscape much more accurately, with most slopes being close to the optimum (i.e., 1; fig. 2). The high quality of long-read based assemblies was also observed with different assembly algorithms (supplementary fig. 3) and with unpolished assemblies (supplementary fig. 2). In agreement with previous work, we thus conclude that long-read based assemblies are well suited for genomic analyses of TEs (Ellison and Cao, 2020). Short-read based assemblies, on the other hand, suffer from few contiguously assembled piRNA clusters and a poor representation of the TE landscape.

**Figure 2:**
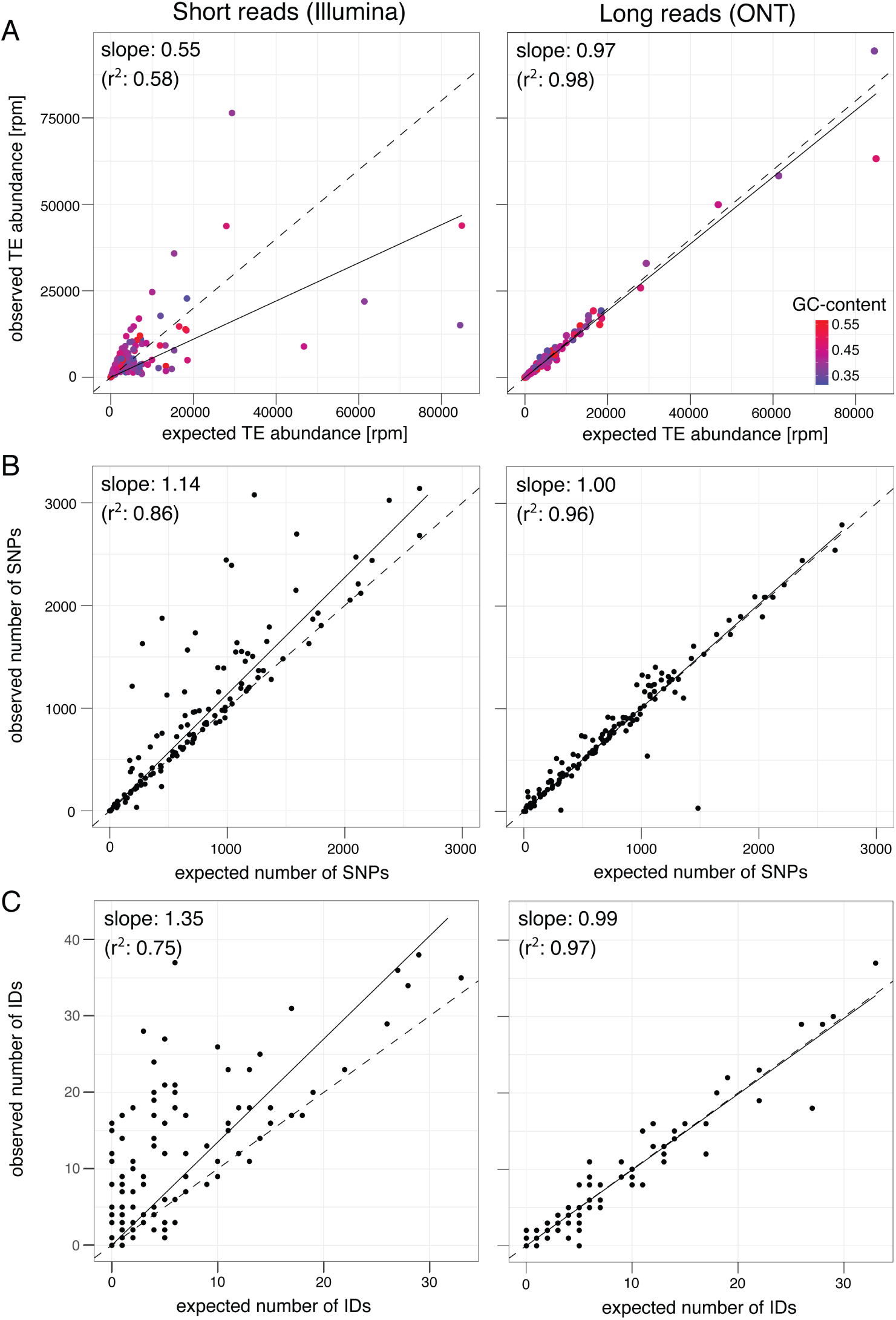
Three novel quality metrics estimate how well an assembly captures the TE landscape of an organism, i.e., for each TE family (black or colored dots), the abundance (A), number of SNPs (B), and internal deletions (IDs) (C). Our metrics are based on a comparison between the expected and the observed TE landscape where the Illumina raw reads are used to derive the expected TE landscape and artificial reads generated from an assembly represent the observed TE landscape. By aligning the reads to the consensus sequences of TEs, the expected and the observed abundance of each of the three features (A, B, C) can be quantified. A regression between expected and observed values allows us to summarize the results across all analyzed TE families. The slope of these regressions represent three of our novel quality metrics. Deviations from the expected diagonal (dashed lines) indicate either an over- or under-representation of a feature in the assembly. To illustrate the new metrics, we compared the quality of a short- and long-read based assembly of Canton-S. Note that the long-read based assembly captures the TE landscape more accurately than the short-read based assembly (despite expectations being based on short reads).

In summary, we introduced four novel quality metrics for estimating the suitability of assemblies for TE research. The first quality metric, the CUSCO value, captures the fraction of contiguously assembled piRNA clusters. Computing this metric requires sequences of regions flanking piRNA clusters. This metric may thus solely be used with organisms already having a reference genome. The last three metrics estimate whether an assembly accurately reproduces the TE landscape. Computing these metrics requires Illumina paired-end reads and consensus sequences of TEs. These metrics may thus also be used with organisms not having a reference assembly (as long as a few TE sequences are known). We made the scripts, a manual, and a walk-through for computing our quality metrics available https://sourceforge.net/projects/cuscoquality/.

### Optimizing the assembly strategy

Next, we aimed to identify assembly strategies that enable us to generate a high-quality assembly for the *D. melanogaster* strain Canton-S. At first, we evaluated the performance of three different long-read assemblers, which rely on slightly different algorithms. Miniasm (Li, 2016) uses the overlap among reads to build contiguous sequences. Canu (Koren et al., 2017) utilizes a similar approach as miniasm. However, to reduce the error rate, Canu trims reads and generates consensus sequences of reads prior to the assembly. Wtdbg2 (Ruan and Li, 2020) uses a de-Bruijn graph based assembly algorithm, where k-mers are much larger than for short reads. Long reads usually have high error rates, and assemblies based on these reads may thus also contain appreciable amounts of errors (Sović et al., 2016; Vaser et al., 2017). Following recommendations of previous works (Solares et al., 2018; Chakraborty et al., 2019; Ellison and Cao, 2020), we aimed to reduce the error rate by polishing the assembly with Racon (long reads) (Vaser et al., 2017) and Pilon (short reads) (Walker et al., 2014). Polishing algorithms align reads to an assembly and infer the consensus sequence (Vaser et al., 2017; Walker et al., 2014). Initially, we were concerned that this procedure could eliminate polymorphisms from TE sequences, such that the number of SNPs and IDs of TEs may be underestimated in polished assemblies. Interestingly, we found that polished assemblies capture the TE landscape slightly more accurately than unpolished assemblies (supplementary fig. 2,5). Polishing thus enhances the suitability of assemblies for TE research. We performed one to three round of polishing with Racon and Pilon, where the optimal number of iterations was selected based on the maximally attained BUSCO values (supplementary table 3).

To investigate the influence of coverage on assembly quality, we evaluated the performance of each assembler with several different coverages. Reads were randomly subsampled to coverages ranging from 20-150x (fig. 3). Note that a minimum coverage of 20x was required for Canu and wtdbg2. To assess the quality of the assemblies, we used our novel TE-centered quality metrics and classical metrics (NG50, BUSCO, and assembly length) (fig. 3; supplementary table 4). We found that the quality of the assemblies depended on both the coverage and the assembler (fig. 3). Interestingly, the best assemblies were not necessarily obtained when all reads were used (fig. 3). For example, Canu yielded the largest NG50 with a coverage of 100x and miniasm the best representation of TEs at a coverage of 50x. Based on our TE-centered quality metrics (abundance, SNPs, IDs, and CUSCO), Canu and miniasm outperformed wtdbg2 at all evaluated coverages (fig. 3). At most coverages, Canu captured the TE abundance more accurately than miniasm and wtdbg2 (fig. 3). Furthermore, assemblies generated with Canu mostly had the highest CUSCO values (fig. 3), where up to 80% of the piRNA clusters were contiguously assembled with coverages ranging from 100x to 150x. While wtdbg2 yielded the highest NG50 values, it also generated the shortest assemblies (fig. 3). The Canu assemblies were the largest at most coverages, and the NG50 values were intermediate (fig. 3). BUSCO values were very similar among the evaluated coverages and assemblers, suggesting that BUSCO is of limited use for estimating the suitability of assemblies for TE research (supplementary table 4). Overall, we conclude that Canu yields the best assemblies from the perspective of TE biology: most piRNA clusters were contiguously assembled, and the TE landscape was most accurately captured (fig. 3). Thus, for the remainder of this manuscript, we relied on assemblies generated with Canu. Interestingly, the best assemblies with Canu were not obtained when the full data set was used (150x), but rather, when a randomly chosen subset of the reads was used (e.g., 100x; 3; NG50 and IDs). When reads are randomly sampled, large portions of the data will not be used for the assembly. These unused data may, however, still contain long reads that could be useful for improving the quality of assemblies, e.g., by bridging gaps between contigs. Thus, we next asked if the assembly quality could be further enhanced by sampling the longest reads instead of a random subset. To test this, we sampled subsets of the longest reads with coverages ranging from 20x to 150x (fig. 4). The mean read length of these subsets ranged from 25, 051 bp with 20x coverage to 7, 146bp with 150x coverage (fig. 4A). Canu assemblies based on the longest reads usually have higher NG50 values than assemblies based on random reads (fig. 4B). The largest NG50 values were again obtained when a coverage of 100x was used (fig. 4B). Note that 150x coverage corresponds to the full dataset. Hence, no differences between assemblies based on random reads and the longest reads are expected at this coverage (fig. 4B,C).

**Figure 3:**
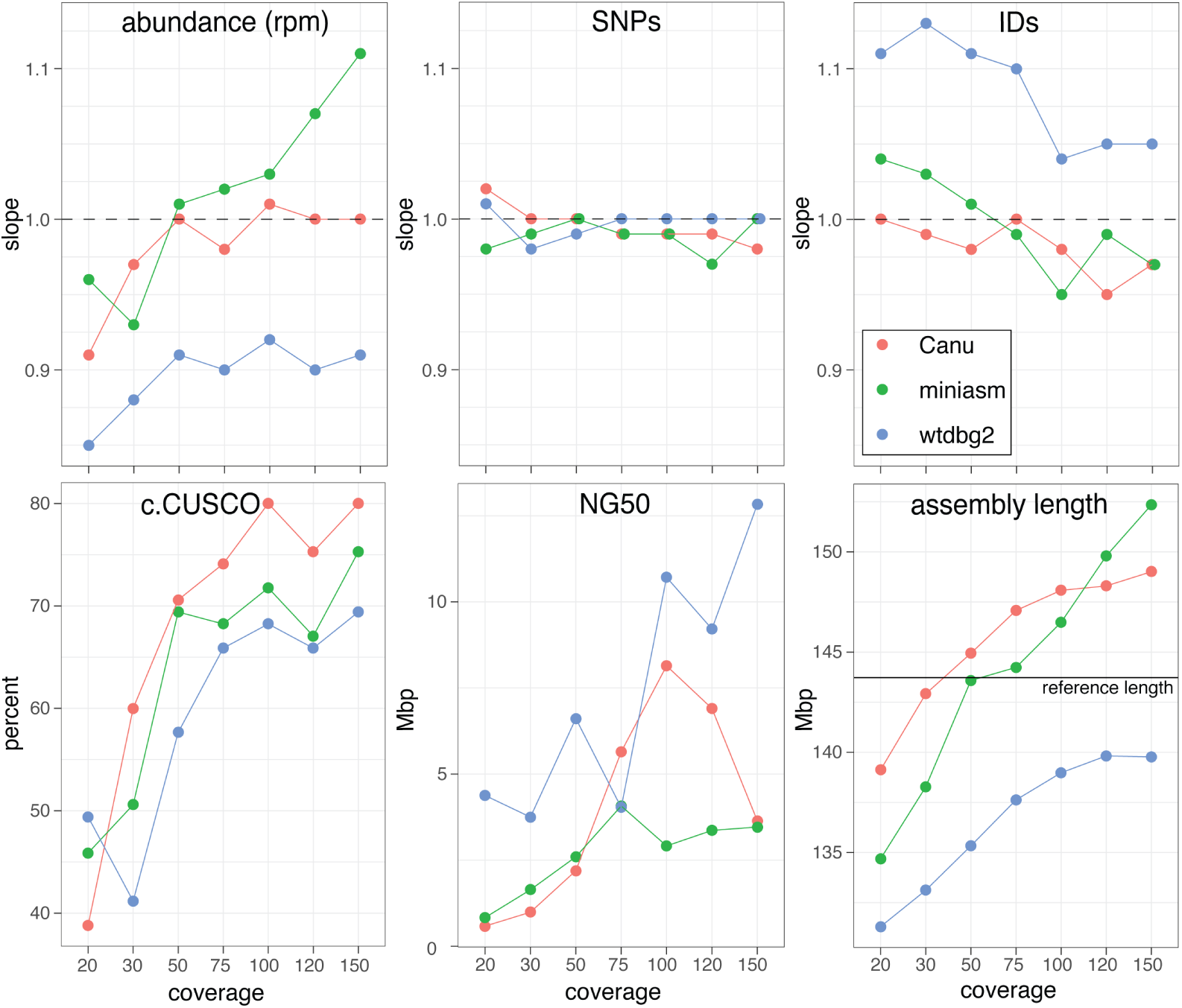
Influence of the assembly algorithm (Canu, miniasm, wtdbg2) and the coverage on the quality of assemblies. Results are shown for our novel TE-centered quality metrics (CUSCO, abundance, SNPs, IDs) as well as classic quality metrics (NG50, BUSCO). Dashed lines indicate optimal performance.

**Figure 4:**
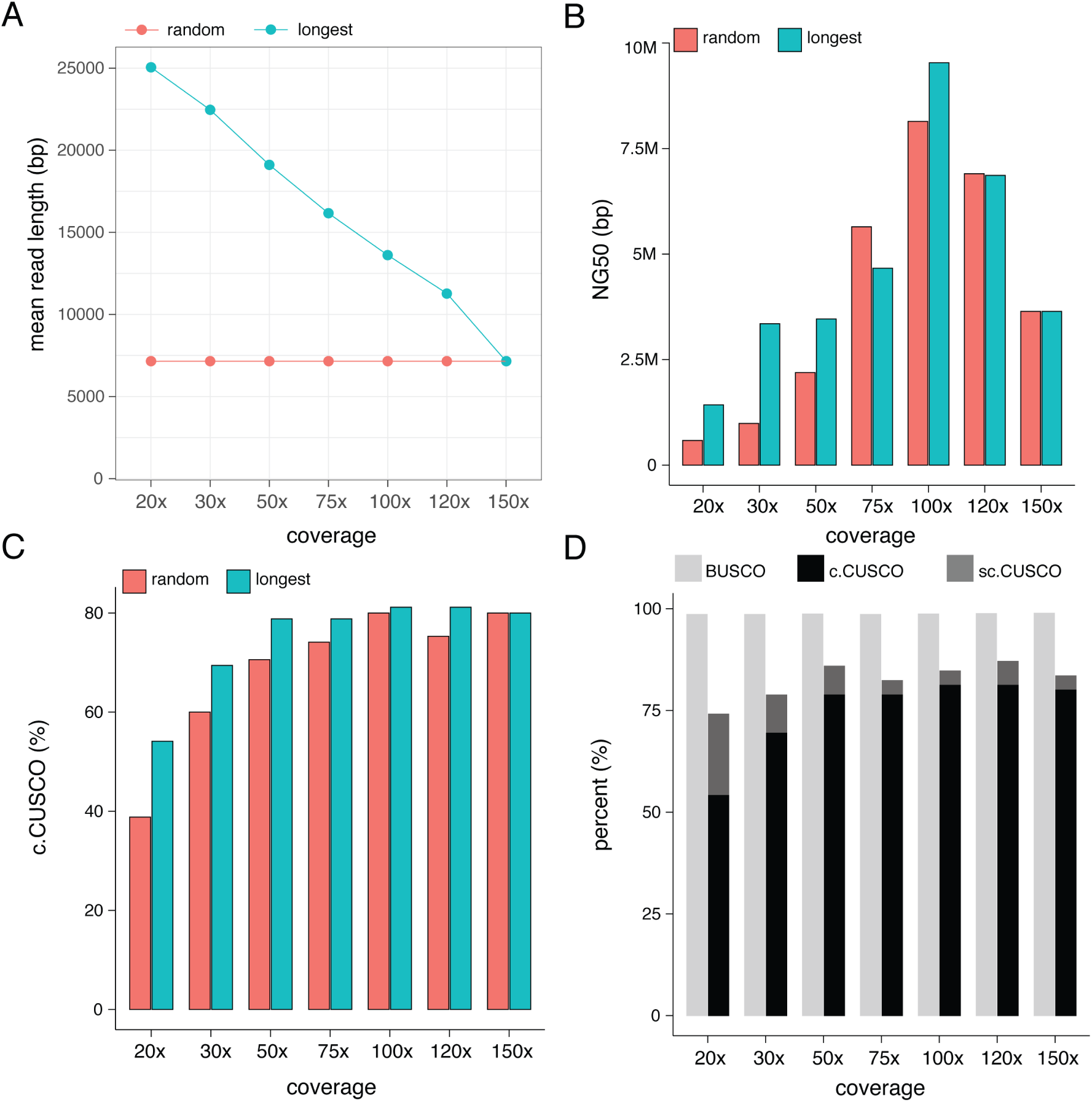
Assemblies with Canu have the highest quality when solely a subset of the longest reads is used. Assemblies were either based on a random subset of the reads (random) or on the longest reads (longest). A) Mean read length of the investigated subsets of reads. B) NG50 values of assemblies generated with different subsets of reads. C) contig-CUSCO values of assemblies generated with different subsets of reads. D) CUSCO and BUSCO values of assemblies generated with different subsets of the longest reads. Scaffolding was based on Hi-C data.

Interestingly, CUSCO values were consistently highest for assemblies generated with the longest reads (fig. 4C), which highlights the importance of long reads for assembling piRNA clusters. The three TE landscape metrics (abundance, SNPs, IDs) revealed little differences between assemblies generated with random reads and the longest reads (supplementary fig. 4).

Finally, we were interested in whether CUSCO values could be further improved by using *de-novo* scaffolding with Hi-C data (fig. 4D). Scaffolding algorithms link contigs into longer sequences based on diverse information such as genetic maps, optical maps or the conformation of chromosomes (Rice and Green, 2018). One widely used approach for scaffolding, Hi-C, relies on the three-dimensional organization of chromosomes (Lieberman-Aiden et al., 2009). With Hi-C, chromatin interactions may be identified by sequencing fragments that were physically in close proximity (Rice and Green, 2018; Sedlazeck et al., 2018a). Since chromatin interactions are most often observed among neighboring sites within chromosomes, Hi-C data can also be used for scaffolding (Rice and Green, 2018; Sedlazeck et al., 2018a).

Since scaffolds usually contain gaps of unknown size between the contigs (mostly indicated by 100 ‘N’ characters), we calculated the scaffold-CUSCO (fig. 4D). As expected, scaffolding with Hi-C data substantially increased the NG50 value of the assemblies (145 − 1033%; supplementary fig. 5). Interestingly, despite this substantial increase in NG50 values, scaffolding with Hi-C data only led to a moderate improvement of the CUSCO value (3.5 − 20%; fig. 4D). The gain in CUSCO value was most pronounced at low coverages, where the long-read based assemblies had the smallest CUSCO values. Although Hi-C data may substantially improve the contiguity of assemblies, they are therefore only of moderate use for assembling piRNA clusters: only a few additional piRNA clusters were assembled, but many of these newly assembled clusters can not be used for TE research as they contain gaps of unknown size (see below). As expected, other quality metrics, such as BUSCO and the three TE landscape metrics, were not influenced by Hi-C based scaffolding (supplementary table 5).

In summary, we found that our novel metrics are useful for assessing the quality of assemblies. Especially CUSCO turned out to be a sensitive metric that identified quality differences not found by other metrics. With our optimized assembly strategy, up to 81% of the piRNA clusters could be contiguously assembled in *D. melanogaster*. Especially assemblies based on Canu and a subset of the longest reads (100x coverage) had a high quality. Finally, we found that Hi-C data were of limited use for assembling piRNA clusters.

### Influence of segregating polymorphisms on assembly quality

Based on Canton-S, we showed that long reads enabled us to generate high-quality assemblies for TE research. However, Canton-S is highly isogenic, with few segregating polymorphisms (fig. 5A). Generating such highly isogenic strains is laborious and time-consuming, requiring many generations of inbreeding or sophisticated crossing schemes (Brizuela et al., 1994). It would be beneficial if high-quality assemblies could be generated for less isogenic strains. Being able to assemble outbred strains caught from natural populations directly could, for example, substantially simplify investigating the population dynamics of TEs. We used the *D. melanogaster* strain Pi2 to assess the influence of segregating polymorphism on assembly quality. This strain is frequently used in TE research, for example, to assess the extent of P-element (a DNA transposon) induced infertility in females (O’Hare and Rubin, 1983; O’hare et al., 1992; Srivastav et al., 2019) and has substantial numbers of segregating SNPs on several chromosomes. (fig. 5A). We first generated a high-quality data set for Pi2: 199x ONT long reads (mean read length ≈ 8kb), 40x of Illumina PE data, and 260x coverage Hi-C data (supplementary table 1). We generated the Pi2 assembly following our previously established strategy: 100x of the longest ONT reads, with mean read length 19, 219 bp, were assembled with Canu, assemblies were subject to multiple rounds of polishing, and Hi-C data were used for scaffolding (supplementary table 5). The Pi2 assembly has a similar BUSCO value to the Canton-S assembly (supplementary table 5). Interestingly, the CUSCO value of the Pi2 assembly is higher than of the Canton-S assembly (supplementary table 5). The TE landscape is accurately reproduced in the Pi2 assembly, similarly to in the Canton-S assembly (supplementary table 5). Solely based on our TE centered quality metrics, the assemblies of Canton-S and Pi2 are thus of similar quality. We, however, noticed that the Pi2 assembly is substantially larger than the Canton-S assembly (13% larger, supplementary table 5). This difference in assembly size may be due to the polymorphisms segregating in Pi2, where the assembly algorithm could have generated several contigs (e.g., a contig for each homologous chromosome) for polymorphic regions (Pryszcz and Gabaldón, 2016). Such an unequal representation of genomic regions could lead to problems and hard to detect biases during the downstream analyses of assemblies (Pryszcz and Gabaldón, 2016), including the genomic analyses of TEs. To test whether our Pi2 assembly suffers from this problem, we sliced assemblies into non-overlapping fragments of 1kb, aligned them to the reference sequence, and calculated the average coverage for 100 kb windows (fig. 5A). Uniquely assembled regions will have a coverage of 1, whereas regions assembled multiple times will have a coverage *>* 1 and regions missing from an assembly a coverage *<* 1. We observed many multiple-assembled regions for the Pi2 assembly (fig. 5B). These redundant contigs largely overlap with polymorphic regions (fig. 5C). We thus conclude that segregating polymorphism in Pi2 led to redundant contigs, especially in regions with many segregating polymorphisms. These redundant contigs likely account for the large assembly size of Pi2 (supplementary table 5). Interestingly, we did not observe any redundant assemblies of piRNA clusters for Pi2 (we tested if both sequences flanking piRNA clusters map to multiple contigs). This absence of redundant clusters is likely due to the fact that the vast majority of the piRNA clusters lie in regions with few segregating polymorphisms in Pi2 (supplementary fig. 6).

**Figure 5:**
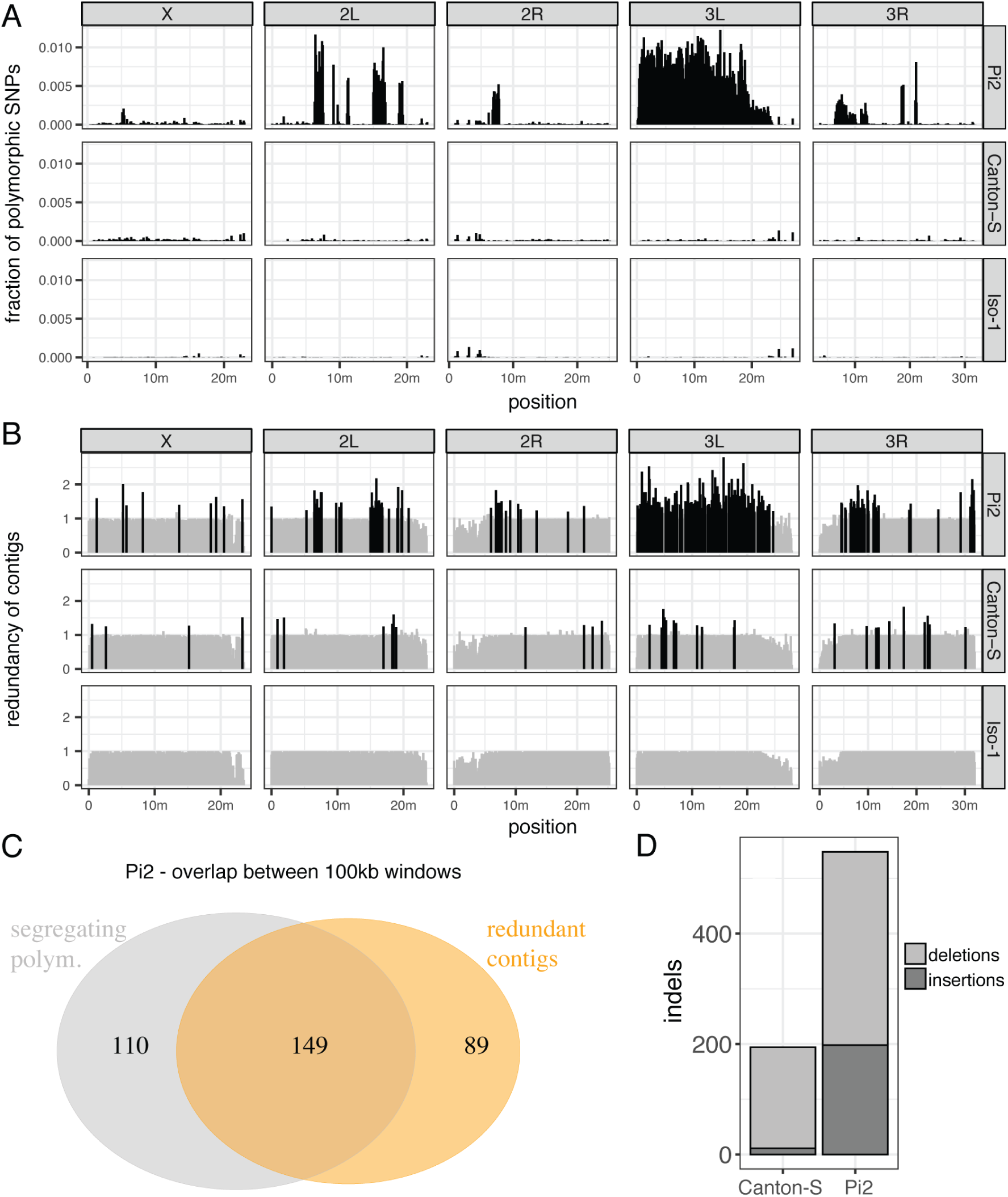
Segregating polymorphisms may lead to redundant contigs. A) The fraction of segregating SNPs for the *D. melanogaster* strains Canton-S and Pi2. The results are shown for 100kb windows. The highly isogenic strain Iso-1 is included as a reference. B) Origin of redundant contigs. Non-overlapping 1kb subsequences of an assembly were aligned to the reference. The average coverage per 100kb window is shown. Coverages *>* 1.2 indicate redundant contigs (shown in black). C) Redundant contigs are mostly found for windows with segregating polymorphism. D) Number of large indels (*≥* 1kb) in the assemblies of Canton-S and Pi2.

It is possible that redundant contigs are especially abundant for assemblies generated with Canu. To test this, we assembled Pi2 also with miniasm and wtdbg2 using 100x coverage of the longest reads. Indeed, miniasm and wtdbg2 generated fewer redundant contigs than Canu (supplementary fig. 7), which confirms that Canu has difficulties assembling polymorphic regions. However, the contig-CUSCO values of assemblies generated with wtdbg2 (57.7) and miniasm (74.12) were much lower than for the Canu assembly (83.5; after polishing and before scaffolding). Hence the evaluated assembly algorithms exhibit a trade-off between the number of contiguously assembled piRNA clusters (Canu is better) and the number of redundant contigs (wtdbg2 and miniasm2 are better).

Polymorphic regions can be a problem for genomic analyses of TEs since heterozygous TEs may be overlooked. This could lead to erroneous conclusions during the downstream analysis, such as the absence of a TE in piRNA clusters, when it is, for example, present in heterozygous form (see the example shown below). We thus looked if we find structural variants (SVs) in our assemblies using Sniffles, which scans long reads aligned to an assembly for SVs (Sedlazeck et al., 2018b). In total, we identified 194 large indels (*>*1kb) for our Canton-S assembly and 548 for our Pi2 assembly (fig. 5D). Deletions (e.g., heterozygous SVs present in the assembly) were more abundant than insertions for both assemblies (fig. 5D). A blast search revealed that 7.7% and 39.1% of the SVs in Canton-S and Pi2, respectively, were due to TEs.

We conclude that an assembly of strains with segregating polymorphisms is challenging and may lead to misleading conclusions. The best assemblies for TE research are obtained when highly isogenic strains are used.

### Polymorphisms in piRNA cluster

Finally, we were interested in whether our high-quality assemblies could be used to gain novel insights into TE biology. It is widely assumed that TE invasions are stopped when a member of the TE family jumps into a piRNA cluster, which triggers the production of piRNAs that repress the TE (the trap model). If this model is correct, we expect to find polymorphic TE insertions in piRNA clusters (Kelleher et al., 2018; Kofler, 2019). To test this hypothesis, we compared the piRNA clusters among our assemblies and the reference genome. In addition to the piRNA clusters of Canton-S, we also used the clusters of Pi2 as the clusters of this strains were non-redundantly assembled (see above). Presence/absence polymorphism of TE insertions within clusters could lead to vast size differences of clusters among strains. Thus, we first investigated the length of the piRNA clusters (i.e., the distance between the two sequences flanking each cluster). Interestingly, the cluster lengths of both, Canton-S and Pi2, are very similar to the reference genome (release 6; fig. 6A). Solely 14 clusters in Pi2 and 16 clusters in Canton-S deviated in length by more than 20% from the length of the clusters in the reference genome. Some of this size variation (7 in Pi2 and 2 in Canton-S) was due to clusters with a gap in the assembly (recognized by several ‘N’ characters; fig. 6A, colored dots). Despite the similar length of the clusters, the composition of clusters may vary substantially among strains. To test this, we carefully investigated the cluster 42AB, one of the largest clusters in *D. melanogaster*. 42AB is a germline-specific piRNA cluster with a size of 248 kb. It is located on the chromosome arm 2R between the genes *jing* and *Pld*. We annotated TEs with RepeatMasker (Smit et al., 2015) and identified sequence similarity between our assemblies and the reference genome with Blast (fig. 6B; (Altschul et al., 1990)). Most TE insertions are shared between the three strains. Even large synteny blocks, frequently involving several TE insertions, can be found (fig. 6B). Nevertheless, we also found differences among the three strains (fig. 6B). Most notably, a 26kb region - involving the X-element, GATE, Max-element, and rover - was duplicated in Pi2 (fig. 6B). Relative to the reference genome, we also found a substantial number of TE presence/absence polymorphism in both strains (4 in Pi2 and 12 in Canton-S; fig. 6B). Interestingly, most of these differences were due to TEs having a low divergence from the consensus sequence of the TE (*<*1%; fig. 6B), which suggests that these polymorphisms are due to novel TE insertions into 42AB. Apart from Chimpo, which was identified using RepBase (Bao et al., 2015), all TEs identified in the cluster 42AB were present in the consensus sequences of TEs in *D. melanogaster* (v10.01; (Quesneville et al., 2005)).

**Figure 6:**
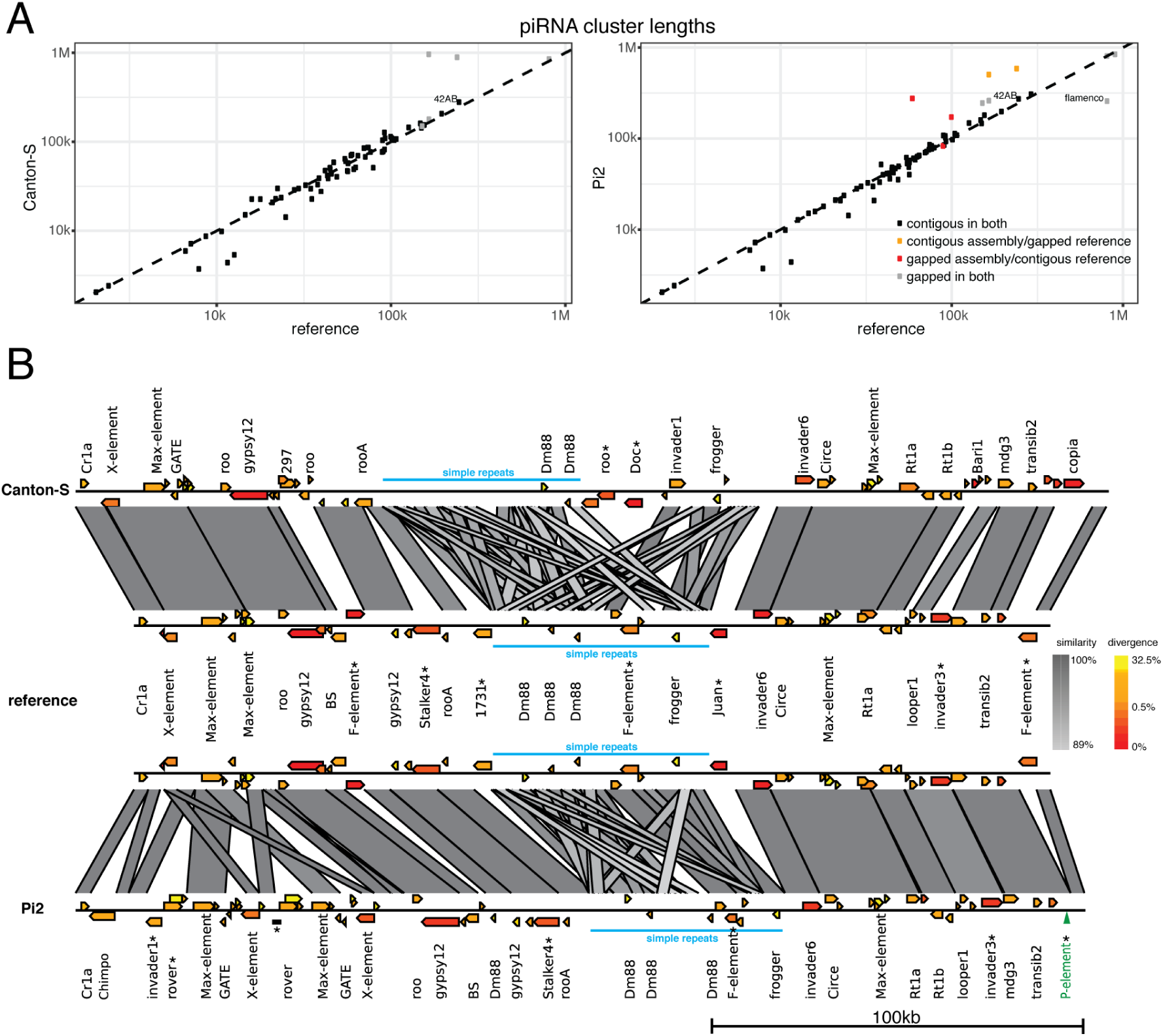
piRNA clusters have presence/absence polymorphisms of TE insertions. A) The length of piRNA clusters in our assemblies is similar to the length of the clusters in the reference genome (dashed line). Clusters with assembly gaps (i.e., ‘N’ characters in assembly) are indicated in color. B) The sequence of the cluster 42AB in our assemblies compared to the reference genome. The TE annotation (yellow-red gradient indicates similarity to the consensus sequence of the TE) and sequence similarity to the reference genome (grey gradient indicates the degree of similarity) are shown. PCR validated presence/absence polymorphisms of TEs or SVs are marked with a ‘*’. TE insertions not present in the assembly are shown in green.

We were interested in whether TE insertions could have been missed in our assembly of 42AB (e.g. heterozygous insertions). Using Sniffles, we indeed found one insertion: a P-element insertion at the distal end of 42AB in the strain Pi2 (fig. 6B; green). The P-element is a DNA transposon, that recently invaded *D. melanogaster* populations. The presence of the P-element in 42AB is in agreement with a simulation study which proposed that TEs that recently invaded a novel species should have segregating piRNA cluster insertions among individuals (Kofler, 2019). Furthermore, this also demonstrates that segregating TE insertions may be missed in assemblies and that this could lead to wrong biological conclusions (e.g., that the P-element is not present in piRNA clusters).

Finally, we validated several of the polymorphic TE insertions in piRNA clusters with PCR. In the cluster 42AB we confirmed 12 out of the tested 14 polymorphic TE insertions, including the missing P-element insertion and the large duplication in Pi2 (7 present in Pi2; 2 present in Canton-S; 7 present in Iso-1 of which three are shared with Pi2 supplementary fig. 8; supplementary table 6). In other piRNA clusters, we confirmed 20 out of the 22 tested polymorphic TE insertions (12 present in Pi2; 10 present in Canton-S; supplementary fig. 8; supplementary table 6).

We conclude that our assembly strategy yields complete sequences of many piRNA clusters. Our assemblies are thus suitable for genomic analyses of TEs and may lead to novel insights into the biology of TEs. For example, we found that piRNA clusters carry abundant presence/absence polymorphisms of TEs, which supports the view that piRNA clusters act as traps for invading TEs (Bergman et al., 2006; Kelleher et al., 2018; Kofler, 2019).

### Finalizing assemblies

To provide chromosome-scale assemblies of Pi2 and Canton-S to the community, we manually broke up misassemblies (supplementary fig. 9) and performed reference-based scaffolding with RaGOO (Alonge et al., 2019). Reference-based scaffolding raised the scaffold-CUSCO to 95.3 for Canton-S and to 97.7 for Pi2 but had little effect on other quality metrics (supplementary table 5). Note that most piRNA clusters assembled by scaffolding have gaps (several ‘N’ characters) and are thus solely of limited use for TE research. An overview of the quality of the final assembly, including the quality at the different assembly steps, can be found in supplementary table 5. The assemblies of Canton-S and Pi2 are available at NCBI (PRJEB37493)

## Discussion

Here we showed that long-read sequencing technologies enable us to generate high-quality assemblies suitable for genomic analyses of TEs. With an optimized assembly strategy, more than 80% of the piRNA clusters may be contiguously assembled, and the TE landscape of an organism may be accurately reproduced.

### Novel quality metrics

Since current metrics of assembly quality ignore TEs, it was first necessary to introduce four novel quality metrics. The first metric, the CUSCO value, estimates the fraction of contiguously assembled piRNA clusters based on an alignment of unique sequences flanking the clusters. For several reasons, the contiguity of piRNA clusters is a useful metric for assessing the quality of assemblies. First, piRNA clusters are of exceptional interest to TE biology (Brennecke et al., 2007; Yamanaka et al., 2014). Hence, contiguous assemblies of these regions are a valuable asset for research. Second, the metric allows us to differentiate between assemblies of very different qualities, as the difficulty of assembling a piRNA cluster varies substantially among the clusters. Long clusters may, for example, be much more challenging to assemble than short ones. This broad range of CUSCO values is demonstrated by our assemblies of Canton-S, where the contig-CUSCO ranges from 5.88% (short reads) to 81.18% (long reads). Yet, even further improvements of the assembly quality could be quantified, as illustrated by the remaining gap to the maximum value (97.65% for reference 6 of *D. melanogaster*). For this reason, we argue that CUSCO is a sensitive quality metric capable of differentiating among assemblies of diverse qualities, even when assemblies have a similar quality according to other metrics such as BUSCO (fig. 3; supplementary table 4). Obviously, CUSCO can solely be used for species with piRNA clusters. Since piRNA clusters were found for many species, ranging from insects to mammals, (Brennecke et al., 2007; Ernst et al., 2017; Yamanaka et al., 2014), the CUSCO metric is, in principle, widely applicable. For species without piRNA clusters, such as plants, regions with similar functions, like the KNOT regions of *Arabidopsis thaliana* (Grob et al., 2014), could be used. Alternatively, flanking sequences could be designed for any set of complex regions. Identification of the sequences flanking piRNA clusters, which is necessary for computing the CUSCO, requires a reference genome. Hence, CUSCO values can solely be estimated for species with a reference genome and an annotation of piRNA clusters. But even for species with a reference genome, it will not be feasible to identify suitable flanking sequences for all piRNA clusters. For example, in *D. melanogaster* 46 of the clusters identified by (Brennecke et al., 2007) were on a pseudoscaffold (chromosome-U) that consists of concatenated contigs of unknown origin (Dos Santos et al., 2015), which makes identification of flanking sequences challenging. Moreover, flanking sequences can not be identified for clusters at terminal ends of contigs/chromosomes. However, for computing CUSCO values, it is not necessary to design flanking sequences for all piRNA clusters, although the resolution of CUSCO will be highest when a majority of the clusters are used. For computing CUSCO values, we assumed that a cluster was contiguously assembled if both sequences flanking the cluster align to the same contig/scaffold. This is based on the assumption that major structural variants, e.g., translocations and inversions, involving piRNA clusters, are rare and that the reference genome is reliable. For computing CUSCO values, we ignored the distance between the flanking sequences and thus the size of the piRNA clusters, since theoretical work suggests that clusters may carry many segregating TE insertions (Kelleher et al., 2018; Kofler, 2019) which could lead to substantial size variation of the clusters among strains. Interestingly, our results did not confirm this expectation: we found that clusters have very similar sizes among the three investigated strains (Canton-S, Pi2, and Iso-1; fig. 6A). At least in *D. melanogaster* it may thus be valid to take the size of piRNA clusters into account by, for example, allowing a maximal difference between the expected and the observed size of piRNA clusters (our CUSCO script supports providing such a parameter). Finally, it is important to distinguish between contig- and scaffold-CUSCO values. The contig-CUSCO is biologically more relevant as it solely estimates the fraction of completely assembled piRNA clusters, whereas the scaffold-CUSCO also considers clusters with gaps of unknown size. Scaffolding algorithms, which introduce gaps between neighboring contigs, have been used to generate most publicly available assemblies (supplementary fig. 10). To gain a complete picture of assembly quality, we thus recommend computing both CUSCO values (supported by our CUSCO script).

However, once sequences flanking piRNA clusters were identified, CUSCO values can be readily computed for many different assemblies. To illustrate this, we estimated CUSCO values for several publicly available assemblies of different *D. melanogaster* strains (supplementary fig. 10; (McCoy et al., 2014; Vicoso and Bachtrog, 2015; Anreiter et al., 2017; Singhal et al., 2017; Chakraborty et al., 2019; Ellison and Cao, 2020)). Assemblies based on ONT and PacBio long reads have a similar quality than our assemblies of Pi2 and Canton-S (supplementary fig. 10). Interestingly, an assembly based on synthetic long reads (Illumina TruSeq) has a much lower CUSCO value than the other long-read based assemblies (supplementary fig 10; (McCoy et al., 2014; Ellison and Cao, 2020; Chakraborty et al., 2019)). As expected, assemblies based on Illumina short reads have much smaller CUSCO values than assemblies based on long reads (supplementary fig. 10).

Computing CUSCO values requires sequences flanking piRNA clusters. These sequences need to be identified for each species separately. We made the sequences flanking piRNA clusters of *D. melanogaster* publicly available https://sourceforge.net/projects/cuscoquality/ and invite the community to contribute flanking sequences for any species of interest to our repository.

With three additional metrics, we estimate whether an assembly accurately captures the TE landscape (abundance, SNPs, and IDs) of an organism. Unfortunately, the real TE landscape is not known for any organism. However, we argue that Illumina raw reads may be used to derive a useful approximation of the expected TE landscape. Assuming that reads are more or less randomly distributed over the genome, and that sequencing errors are largely random within reads, this assumption should mostly be valid. Sequencing errors can be largely eliminated from the analysis by using a minimum allele frequency (Kofler et al., 2011a). Here, we used a minimum allele frequency of 2% for SNPs and IDs. In case more stringent criteria are required, a higher threshold may be used. The coverage will fluctuate over the genome, which could affect estimates of TE abundance. Especially the GC-bias, where regions with a high GC content have an elevated coverage (Minoche et al., 2011), could lead to overestimating the expected abundance of TEs with a high GC content. However, since we sum the average coverage over many different insertions of a TE family, with insertion sites in diverse genomic backgrounds (with varying GC contents), the influence of the GC-bias and of stochastic coverage fluctuations should be minimized by our approach. Furthermore, since we rely on the slope between the expected and the observed TE abundance, which is based on many TE families with different GC contents, the influence of the GC bias should be further reduced. In agreement with this, we did not find any correlation between GC-content and TE abundance in the raw reads (supplementary fig. 11). Furthermore, we solely found a small but non-significant difference in GC-content among TEs that are well-represented in genomes compared to TEs that are not well-represented (supplementary fig. 11). Finally, it is reassuring that an assembly based on long reads captures the expected TE landscape more accurately than an assembly based on the short reads which have been used to derive the expected TE landscape (fig. 2).

Apart from sequencing biases, also biases occurring during data analysis, such as mapping and quantifying of reads may occur. Since we use the same pipeline for the raw reads (expectations) and the artificial reads derived from an assembly (observations), these biases should largely be eliminated. However, to minimize such methodological biases, it is important that the raw and artificial reads have the same lengths and that the artificial reads are uniformly distributed over the genome. Taken together, we think that Illumina raw reads provide a useful approximation of the expected TE landscape. A comparison to the observed TE landscape allows us to assess whether the TE abundance and diversity are substantially over- or under-represented in an assembly. Computing the TE landscape metrics requires Illumina short reads for an organism (which are often generated for the polishing of assemblies anyway) and consensus sequences of TEs. These metrics may thus be used for model and non-model organisms (assuming some TE sequences are known).

Our novel quality metrics may not only be used to compare the quality of available assemblies but may also serve as a guide during the assembly procedure, for example, to identify the most suitable assembly algorithm. Our metrics should thus help to generate and to identify genomes suitable for genomic analyses of TEs. Nevertheless, to gain a comprehensive picture of assembly quality, it will be important also to consider other metrics such as NG50, BUSCO, and the total size of assemblies (especially when the genome size is known).

### Assembly strategy

We showed that high-quality genomes suitable for genomic analyses of TEs can be obtained if: i) the sequenced strains are isogenic, ii) sequencing is based on long reads, iii) suitable assemblers, such as Canu are used with an optimized coverage and read length, iv) assemblies are polished using short and long reads and v) some scaffolding approach is used. Unfortunately, the time-consuming isogenization of strains still seems necessary as it helps to avoid redundant contigs (fig. 5). We also discovered a trade-off between some currently available assembly algorithms: Canu yields the best assemblies for TE research, but generates many redundant contigs, while the other evaluated assemblers generate fewer redundant contigs but the assemblies are less suitable for TE biology. It is, however, feasible that future algorithms support assemblies of non-isogenic strains. For example, phased assemblers, such as FALCON-phase (Kronenberg et al., 2018), may yield a separate contig for each homologous chromosome. For these algorithms, segregating polymorphism could even be an advantage as polymorphisms may help to distinguish between homologous chromosomes. We also found that long reads allow generating high-quality genomes for TE research. Assemblies generated by any of the two major long-read technologies, ONT and PacBio, have a high quality (supplementary fig. 10). Assemblies based on Illumina short reads and synthetic long reads (Illumina TruSeq) had a lower quality (supplementary fig. 10).

We showed that Canu yields high-quality genomes from the perspective of TE biology: many piRNA clusters are contiguously assembled, and the TE landscape is accurately captured. The high quality of assemblies generated by Canu was also noted in several previous works (Jayakumar and Sakakibara, 2017; de Lannoy et al., 2017; Solares et al., 2018; Wick and Holt, 2019). Interestingly, the best assemblies were obtained when solely a subset of the long reads was used for an assembly with Canu, i.e., 100x coverage with the longest reads. We suspect that this may be related to algorithmic assumptions on the corrected error rate, which is coverage-dependent and governs the overlap among reads (see Canu manual https://canu.readthedocs.io/en/latest/parameter-reference.html). It is, however, important to be aware of the fact that the best assembly is not necessarily obtained when all reads are used, but rather when a subset of the longest reads is used.

Since long reads have a high error rate, polishing of assemblies using long or short reads is usually recommended (Sedlazeck et al., 2018a; Rice and Green, 2018). Interestingly, polishing also enhances the suitability of assemblies for TE research: polishing increased the fraction of contiguously assembled piRNA clusters as well as the representation of TE abundance and diversity (supplementary table 2, 5).

Scaffolding with Hi-C slightly increased the number of assembled piRNA clusters (using scaffold-CUSCO) but had little influence on the representation of the TE landscape (fig. 4; supplementary table 5). The piRNA clusters assembled by scaffolding will largely have gaps and are thus solely of limited use for TE research. Nevertheless, scaffolding approaches will still be useful, since scaffolding enables generating chromosome-sized sequences, which could be important when the genomic context of a TE insertion is relevant (e.g., whether a TE is close to a telomere or a centromere).

Despite our optimized assembly strategy, up to 19% of the piRNA clusters were not contiguously assembled (supplementary table 5, after polishing). Additionally, manual curation of the final assemblies was necessary to avoid misassemblies (supplementary fig. 9). This demonstrates that assembly strategies may still be improved. Especially promising may be further advances in the length of reads (e.g., by further improvements of pores (Deamer et al., 2016)) and in algorithms generating phased assemblies, which could yield a separate contig for each homologous chromosome. Phased assembly algorithms may allow us to use outbred strains and thus to avoid the time-consuming isogenization of strains. Furthermore, these algorithms avoid the central problem of assemblies of diploid organisms, i.e., that two potentially distinct sequences (i.e., the homologous chromosomes) need to be represented as a single one. Phased assembly algorithms could thus prevent wrong conclusions during the downstream analyses of the assemblies, such as the absence of P-element insertions in piRNA clusters (fig. 6).

### Outlook

Due to recent advances in sequencing technology and in assembly algorithms, it is finally feasible to generate high-quality assemblies for TE research. The assembly strategy and the four quality metrics introduced in this work will help to generate high quality-genomes for TE research. Based on our approach, we generated high-quality assemblies of the strains Canton-S and Pi2, which will be useful resources for TE biology and population genomics. High-quality assemblies will enable us to address many important open questions in TE research, such as the genomic distribution of different TE variants, the extent of negative selection acting on different TE variants and the history of TE invasions in a species. Most importantly, however, high-quality assemblies will finally allow us to analyze the composition and evolution of piRNA clusters. For example, one major open question is whether piRNA clusters carry presence/absence polymorphisms as predicted by theoretical work based on the trap model (Kelleher et al., 2018; Kofler, 2019). Our analysis of the cluster 42AB in three different *D. melanogaster* strains showed that such polymorphisms do indeed exist, thus providing experimental support for the trap model (Kelleher et al., 2018; Kofler, 2019).

## Material and Methods

### Sequencing

The *D. melanogaster* strains Canton-S and Pi2 were obtained from the Bloomington Drosophila Stock Center (BDSC) (Canton-S=64349; Pi2=2384). The reference strain, Iso-1, was kindly provided by Dr. K.A. Senti. We performed Oxford Nanopore Sequencing, Illumina paired-end sequencing, and Hi-C for Canton-S and Pi2 (supplementary table 1). The strain Iso-1 was solely sequenced using the Illumina paired-end technology.

High molecular weight DNA for Oxford Nanopore sequencing was extracted from whole bodies of ∼50 female virgin flies using the Phenol-Chloroform extraction protocol described by Maniatis et al. (1982) using slightly elongated incubation times. The DNA was sheared to a mean fragment length of 20-30 kb with Covaris g-TUBEs (Covaris Inc., Woburn, MA, USA). The length of the DNA was measured with a TapeStation (4200; DNA ScreenTape, Agilent Technologies)). Library preparation was performed with an input of 2-5 *µ*g of sheared DNA following the manufacturer’s protocol (kit LSK108; Oxford Nanopore Technologies; Oxford). We used slightly elongated incubation times. About 1-2 *µ*g of the libraries were run for 48-72 hours on MIN106 flowcells. The DNA concentration was measured with a Qubit fluorometer (broad-range DNA assay) (Thermo Fisher Scientific, Waltham, MA, USA), and the quality of the DNA was controlled with NanoDrop (Thermo Fisher Scientific, Waltham, MA, USA).

DNA for Illumina paired-end sequencing was extracted from whole bodies of 20-30 virgin female flies using a salt-extraction protocol (Maniatis et al., 1982). Libraries were prepared with the NEBNext Ultra II DNA Library Prep Kit (New England Biolabs, Ipswich, MA, USA) using 1 *µ*g DNA. Illumina sequencing was performed by the Vienna Biocenter Core Facilities on a HiSeq2500 platform (2×125bp; Illumina, San Diego, CA, USA).

Hi-C was performed following the Phase Genomics Proximo Hi-C animal Kit (Phase Genomics, Seattle, WA). About 40-50 female third instar larvae were sliced with a razor blade to obtain about 80 mg of tissue. Crosslinking and library preparation was performed according to instructions. Sequencing was performed by the Vienna Biocenter Core Facilities NGS on an Illumina HiSeq2500 platform (2×125bp; Illumina, San Diego, CA, USA).

### Assemblies

Basecalling of raw nanopore reads (fast5 format) was performed with either Albacore (v2.3.4; Oxford Nanopore Technologies, Oxford, GB) or guppy (v2.1.3; Oxford Nanopore Technologies, Oxford, GB). Summary statistics, including mean read length and the total output, were calculated with NanoPlot (De Coster et al., 2018) (v1.20.1).

*De-novo* assembly of the nanopore reads was performed with three different tools: Canu (v. 1.7) (Koren et al., 2017), miniasm (v0.3-r179) (Li, 2016), and wtdbg2 (v2.4) (Ruan and Li, 2020). With Canu, raw nanopore reads were corrected and trimmed prior to the assembly (preset –nanopore-corrected). To generate assemblies with miniasm, we first aligned all reads against themselves with minimap2 (Li, 2018) (v2.16-r922) using a preset for nanopore reads (-x ava-ont). We generated the assemblies with miniasm using default settings. The resulting assembly graph files (gfa) were transformed into fasta-files with awk. We launched wtdbg2 with the raw nanopore reads and a nanopore-specific preset (‘preset2’).

Short-read assemblies with Illumina paired-end reads (read length 125 and mean coverage of 30x) were performed with Abyss (v 2.1.5; abyss-pe (Simpson et al., 2009)) using a k-mer size of 96.

Polishing of long-read based assemblies was carried out in two steps. We first used Racon (Vaser et al., 2017) (v 1.2.1) with the raw nanopore reads mapped to the assembly (minimap2; -ax map-ont; v2.16-r922 (Li, 2018)), and then Pilon (Walker et al., 2014) (v 1.22) with Illumina paired-end reads mapped to the assembly (bwa mem). The optimal number of polishing steps was chosen based on the maximally achieved BUSCO values (supplementary table 3).

Scaffolding of contigs was done with Hi-C following the SALSA2 protocol (Ghurye et al., 2019) (v30.Nov.2018). Briefly, Hi-C reads were mapped to the assembly (bwa bwasw), filtered (https://github.com/ArimaGenomics), and duplicates were removed (picard-tools; v2.18.23; https://broadinstitute.github.io/picard/). The mapped reads were then used for scaffolding with SALSA2 using the parameters: diploid mode (-m yes), restriction enzyme sequence (GATC). An assembly graph was provided if available (for miniasm and Canu). Reference-guided scaffolding was performed with RaGOO (v 1.1, Alonge et al. (2019)) based on release 6 of the *D. melanogaster* reference genome (Hoskins et al., 2015).

Random sampling of reads was performed with seqtk (https://github.com/lh3/seqtk) (v1.3-r106). To obtain subsets of the longest reads, we sorted all reads by length and then used the appropriate number of the first reads (i.e. the longest reads). Polishing of assemblies generated with read-subsets was also carried out with these subsets.

To visualize assemblies, we generated dotplots using nucmer (Kurtz et al., 2004) (v3.1). We aligned assemblies to the main chromosome arms (X, 2L, 2R, 3L, 3R and 4) of the *D. melanogaster* reference (‘mumreference’; with parameters -c 1000 -l 100), created coordinate index files using DotPrep.py and visualized genome alignments with dot (https://dnanexus.github.io/dot/).

The final assemblies were based on Canu using 100x of the longest reads. Misassemblies were identified based on Hi-C heatmaps and alignments of the assemblies to the reference genome (dotplots). Hi-C heatmaps were generated with juicer (Durand et al., 2016) (v 1.7.6) using ‘Sau3AI’ as the restriction enzyme. Heatmaps were visualized and analyzed with Juicebox (1.11.08). Potential misassemblies identified in the Hi-C heatmaps were cross-validated with long reads that were aligned to the assemblies. Breaks in the alignment of the long reads were interpreted as support of an assembly error. Contigs with missassemblies were broken with a custom script ‘introduceBreaks2fasta.py’.

### Quality of assemblies

BUSCO (Benchmarking Universal Single-Copy Orthologs) values were based on the *Diptera* dataset (v3.0.2; 2799 genes) (Waterhouse et al., 2017). QUAST (Gurevich et al., 2013) (v5.0.2; quast-lg) was used to compute basic assembly statistics such as NG50 and the total assembly length. As reference, we provided the genome of *D. melanogaster* (release 6).

Computing our TE landscape metrics (abundance of TEs, number of SNPs and indels within TEs) requires artificial reads for an assembly of interest. We generated an artificial read of length 125 bp starting at each position of the assembly (yielding an uniform distribution). The abundance of TEs, as well as the number of SNPs and internal deletions within TEs, were estimated with DeviaTE (Weilguny and Kofler, 2019) (v.0.3.6). DeviaTE was used with both artificial, and the raw Illumina reads. As reference library, we used the consensus sequences of 127 TE families (Quesneville et al., 2005). Solely SNPs and indels with a minimum frequency of 2% were considered. The GC content for each TE was calculated via a custom python script (‘GC-content-calculator.py’).

The CUSCO metric relies on the annotation of piRNA clusters of *D. melanogaster* release 5 (Brennecke et al., 2007; Hoskins et al., 2007). From the 142 annotated piRNA clusters, we excluded the 46 clusters located on chromosome arm U (unassembled fragments) and 10 terminal clusters. For the remaining 86 clusters, we aimed to identify conserved sequences flanking the clusters at both ends. Two piRNA clusters were adjacent to each other (cluster 8 and 9). We designed sequences flanking both clusters. Flanking sequences were further required to align uniquely to release 6 of the *D. melanogaster* reference genome. We were able to design flanking sequences for 85 piRNA clusters. These sequences had a size between 48 to 8, 219 nucleotides. To compute the CUSCO, the flanking sequences were aligned to an assembly using bwa mem (Li and Durbin, 2009). The CUSCO was computed as the fraction of correctly assembled piRNA clusters, where we assumed that a cluster was correctly assembled when both flanking sequences aligned to the same contig/scaffold. We furthermore distinguish between a contig-CUSCO and a scaffold-CUSCO based on the presence of poly-N sequences between the two sequences flanking a piRNA cluster. Poly-N tracts in assemblies were identified using the script ‘find-polyN.py’.

### PCR

PCR reactions were performed at a volume of 20 *µl*, with 0.05 *U/µl* of Firepol polymerase (Solis BioDyne, Tartu, Estonia), 2.5 *mM* MgCl_2_, 200 *µM* dNTPs, 0.2 *µM* primer and 100 *ng/µl* DNA. See supplementary table 6 for all primer pairs. We used a PCR cycler (Bio Rad CFX Connect, Hercules, CA, USA) with the following program: 5 min. denaturation at 94°C; 30 cycles of denaturation (30 sec. at 94°C), annealing (1 min. at 58°C) and elongation (1 min. at 72°C), followed by 10 min. of final extension at 72°C. The PCR products were loaded on a 1% agarose gel and ran with 120 V for 30 min in TBE buffer.

### Data analysis

To identify heterozygous SNPs, we aligned Illumina paired-end reads to release 6 of the *D. melanogaster* genome with bwa mem (v0.7.17-r1188; Li and Durbin (2009)) using default parameters. Reads with a low mapping quality were removed using samtools (v1.7; Li et al. (2009)), a mpileup file was created (samtools), and allele frequency estimates were obtained using mpileup2sync (PoPoolationTE2 (Kofler et al., 2011b)) with the parameters –fastq-type sanger –min-qual 20. The fraction of heterozygous SNPs for windows of 100 kb was computed with a custom script (polymorphicSNPs from sync.py). To account for sequencing errors, we solely classified SNPs with allele frequencies between 0.25 and 0.75 as segregating. Furthermore, a minimum coverage of 10 was required for each site. Windows with insufficient coverage at more than 25% of the sites were excluded. Finally, solely windows with sufficient coverage in all three samples (Pi2, Canton-S, and Iso-1) were retained.

To identify the redundant contigs, we chopped assemblies into non-overlapping fragments of 1kb fragments using a custom script (chopgenome.py) and aligned them to the release 6 of the *D. melanogaster* genome using bwa bwasw with default parameters (v0.7.17-r1188; Li and Durbin (2009)). Ambiguously mapped reads were filtered with samtools (-q 20) and a mpileup file was generated. The mean coverage for 100kb windows was calculated using a custom script (coverage_from_pileup.py).

We use Sniffles (v1.0.7; Sedlazeck et al. (2018b)) to estimate the number of SVs. Such SVs may either be present or absent in the assembly (classified as deletion and insertion, respectively). We first mapped the long reads to assemblies using NGMLR (v0.2.7; Sedlazeck et al. (2018b)) with the parameter -x ont (ONT data as input). SVs were identified with Sniffles using the parameters –report_seq (obtain the sequence of SVs), and –genotype (report allele frequency estimates of SVs). The resulting vcf-file was filtered for SVs with a minimum length of 1kb. To identify SVs caused by TEs, we aligned the sequences of SVs to the consensus sequences of TEs (Quesneville et al., 2005) using blastn.

The composition of piRNA clusters was visualized with easyfig (v2.2.3 08.11.2016; (Sullivan et al., 2011)). Annotations of TEs were obtained with RepeatMasker (open-4.0.7; Smit et al. (2015)) using the parameters: -no_is (skip checking for bacterial insertions), -nolow (skip masking low complexity regions) and *D. melanogaster* TE sequences (Quesneville et al., 2005) or *Drosophila* TE sequences (Bao et al., 2015). Synteny within piRNA clusters among the assemblies was identified with blastn (2.7.1+, Altschul et al. (1990)). To avoid cluttering of the figure, we removed annotations of TEs smaller than 1 kb.

All statistical analysis was done with R (v3.4.3) (R Core Team, 2012) and visualization were performed using the ggplot2 library (Wickham, 2016).

## Availability

Scripts for computing the four quality metrics, including a manual and a walkthrough, are available at https://sourceforge.net/projects/cuscoquality/. The assemblies of Canton-S and Pi2 and the reads are available at NCBI (PRJEB37493). The positions of all contiguously assembled piRNA clusters in Canton-S and Pi2 are available at https://sourceforge.net/projects/cuscoquality/files/publicationdata/piRNA-cluster/ and all scripts used in this work are available at https://sourceforge.net/projects/cuscoquality/files/publicationdata/scripts/.

## Author contributions

RK, FS, and FW conceived this work. FS and OC generated the data. FW performed PCR. FS and FW analyzed the data. RK provided software. RK, FS, and FW wrote the manuscript.

## Acknowledgments

We thank Kirsten Senti for advice and providing the *D. melanogaster* strain Iso-1, Christos Vlachos for sharing scripts, Elisabeth Salbaba for technical support, and all members of the Institute of Population Genetics for feedback and support. This work was supported by the Austrian Science Foundation (FWF) grants P30036-B25 to RK and W1225.

