## supplement for "Generating high quality assemblies for genomic analysis of transposable elements"

March 27, 2020

### 1 Supplementary figures

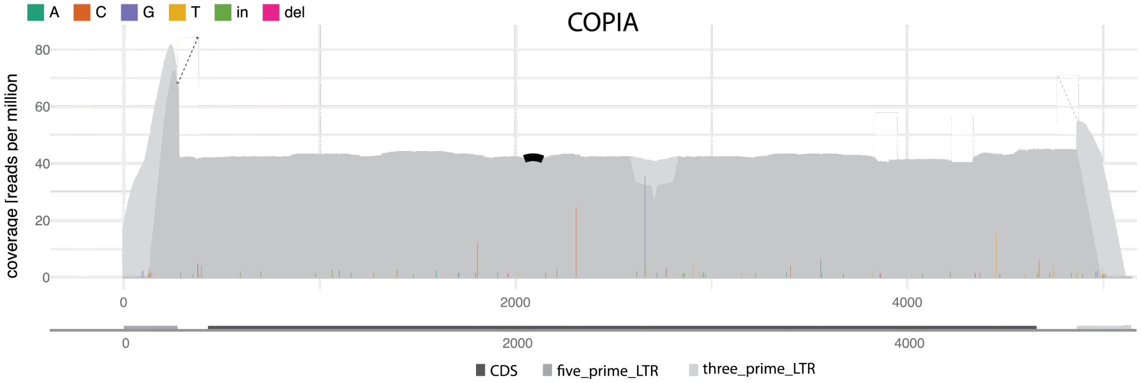


Figure 1: Abundance and diversity for *copia* elements in the *D. melanogaster* strain Canton-S. The coverage (TE abundance in rpm), the position of SNPs (colored lines) and the position of indels (bold arc at the top) are shown. The coverage based on unambiugously (dark grey) and ambiguously (light grey) mapped reads are shown. The plot was generated by DeviaTE (Weilguny and Kofler, 2019) based on Illumina paired end reads mapped to the consensus sequence of *copia* (30 coverage; 2x125bp) and

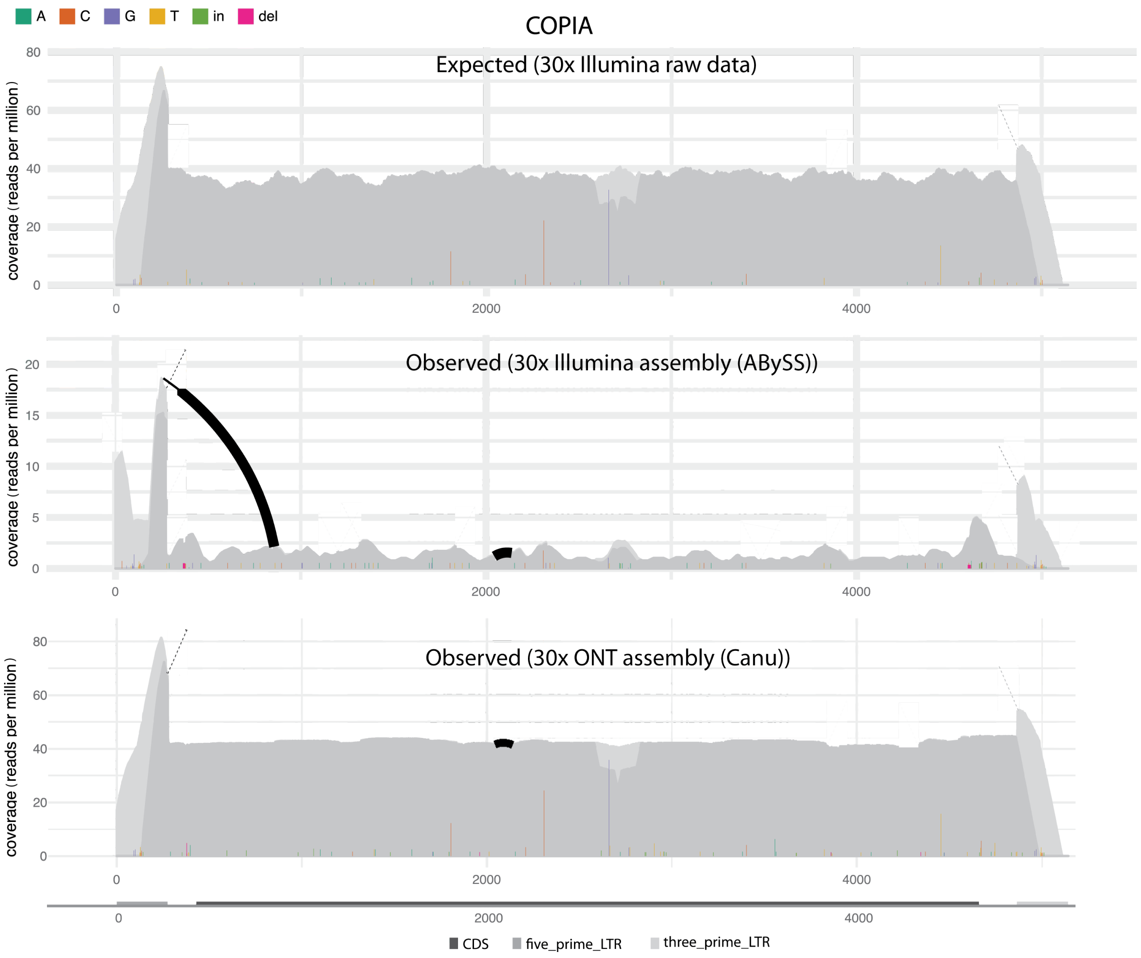


Figure 2: Expected and observed abundance and diversity of *copia* elements in Canton-S. Expected values are based on Illumina raw reads aligned to the consensus sequence of *copia*. Observed values are shown for assemblies based on short (ABySS) and long (Canu) reads. The normalised coverage is shown for ambiguously (light grey) and unambiguously (dark grey) mapped reads. The positions of SNPs (colored lines) and the position of indels (bold arcs) are shown. Note that both, the expected coverage (TE abundance) and diversity (SNPs and indels) of copia, are best reproduced by the long-read based assembly (Canu).

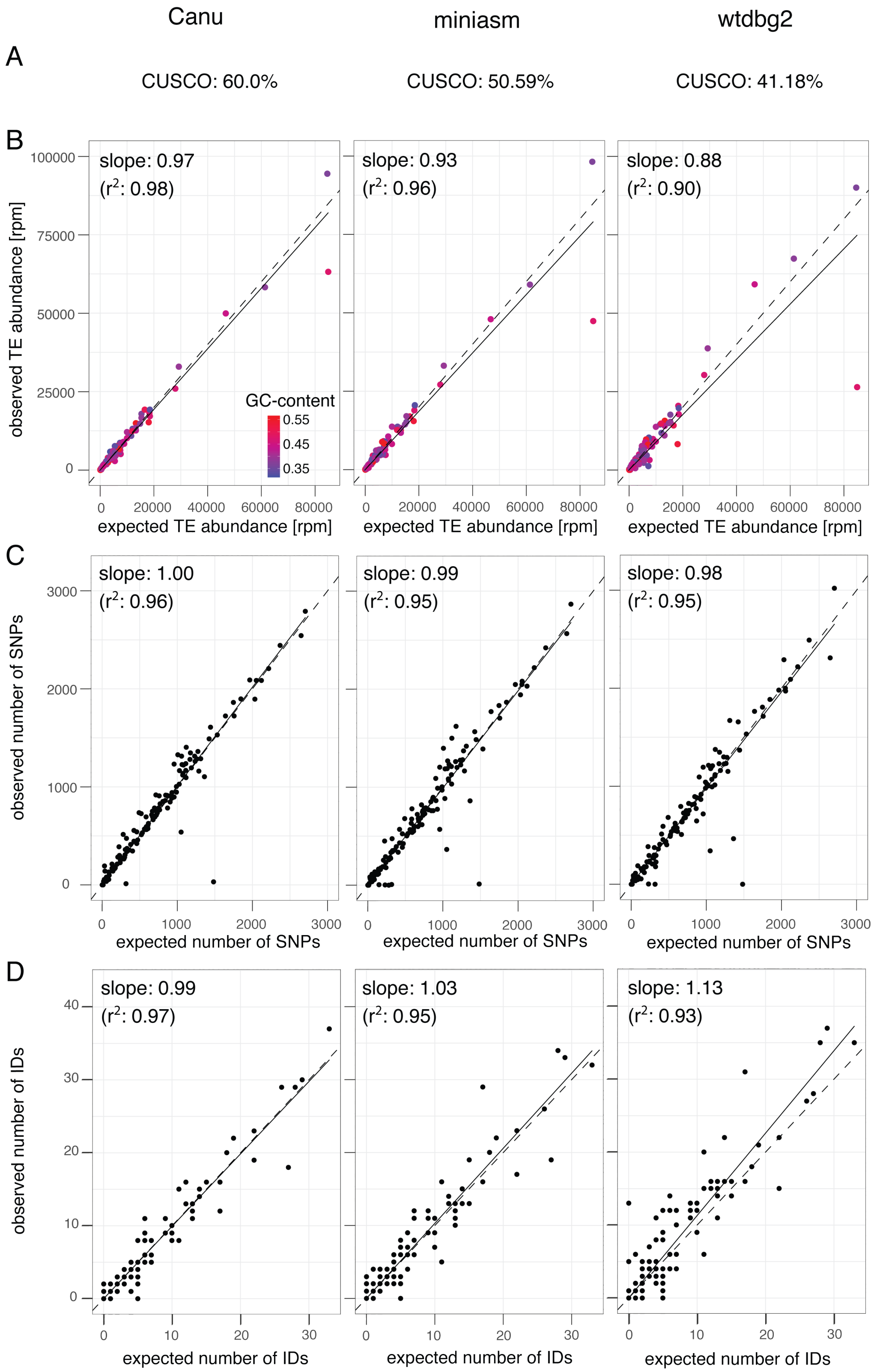


Figure 3: Influence of the assembly algorithm on the quality of assemblies of the *D. melanogaster* strain Canton-S. Assemblies are based on 30x coverage with ONT reads.

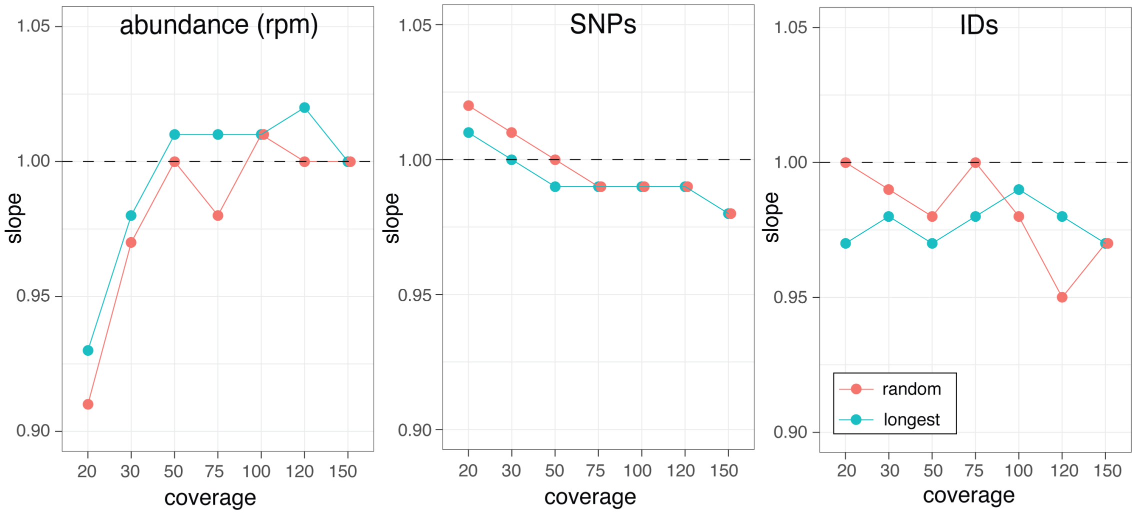


Figure 4: Assembly quality with different subsets of reads. Either random reads (random) or the longest reads (longest) were used to generate assemblies with Canu. The assembly quality is assessed using three of our TE-centered quality metrics (abundance, SNPs, IDs). The dashed lines indicate the optimal representation of TEs.

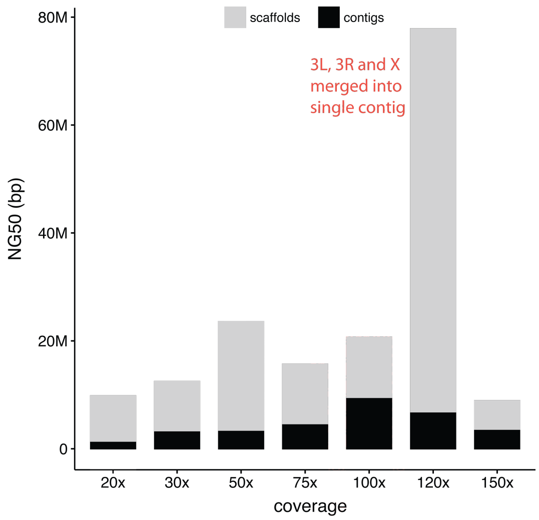


Figure 5: NG50 values of Canton-S assemblies generated with Canu and different subsamples of the longest reads. Values are shown before (contigs) and after Hi-C based scaffolding.

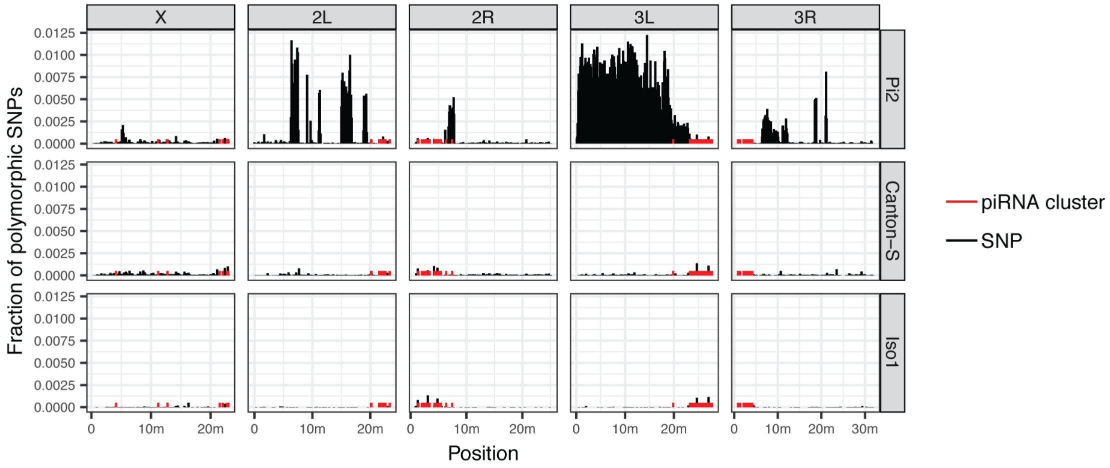


Figure 6: Location of piRNA clusters (red) and of regions with segregating polymorphisms (black) for several *D. melanogaster* strains. Segregating polymorphisms are shown for 100kb windows.

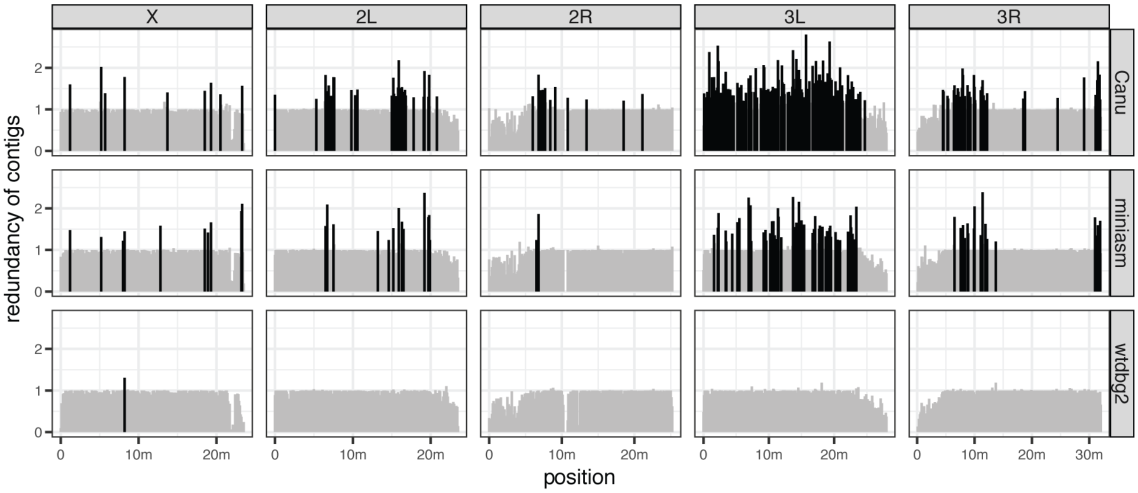


Figure 7: Origin of redundant contigs for assemblies generated by three different algorithm (right panel). Non-overlapping 1kb subsequences of an assembly were aligned to the reference. The average coverage per 100kb window is shown. Coverages > 1.2 indicate redundant contigs (shown in black).

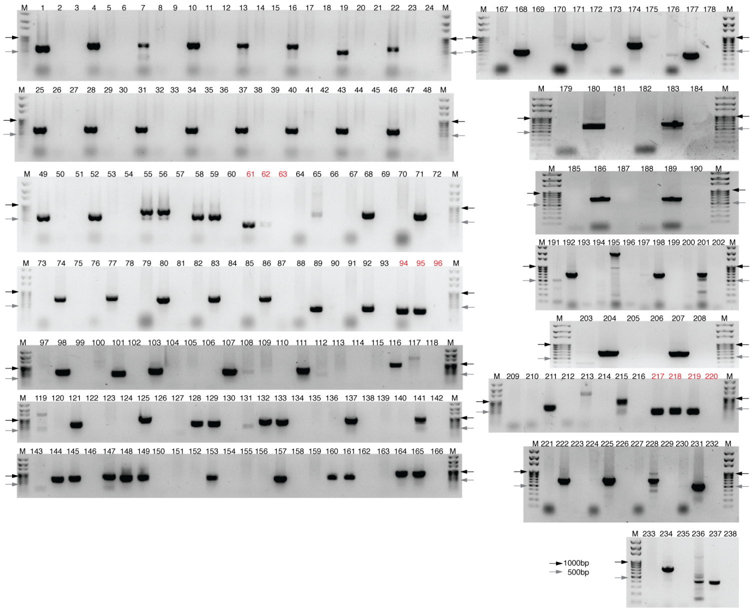


Figure 8: PCR validation of polymorphic TE insertions in piRNA clusters. Numbers above lanes refer to entries in supplementary tables 6 ; Arrows indicate the position of the 1000bp and 500bp size markers. Positive controls (*RpL32*) are labeled in red.

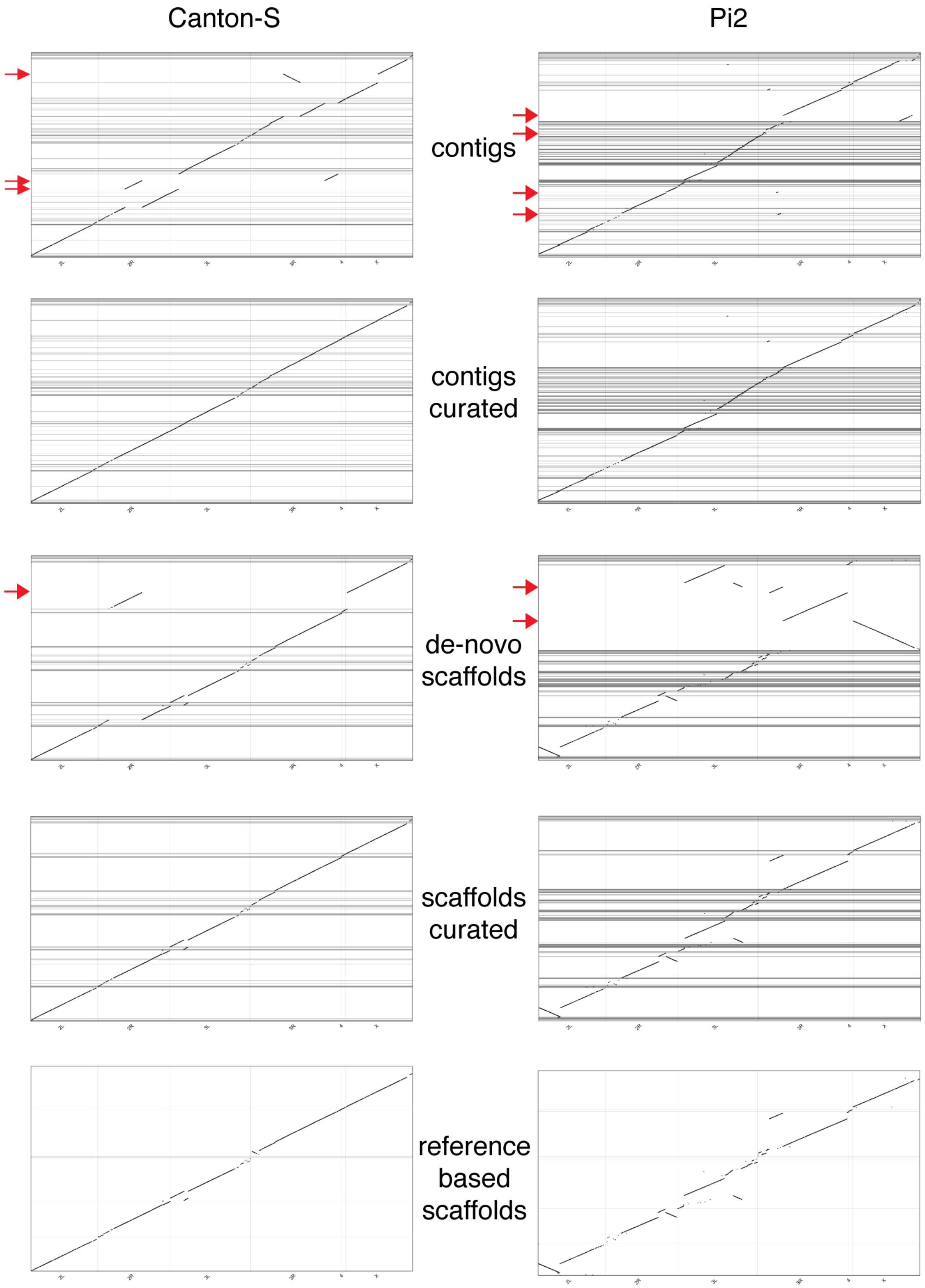


Figure 9: Manual curation steps of the final assemblies of Pi2 and Canton-S. Misassemblies (red arrows) were manually broken up at each step.

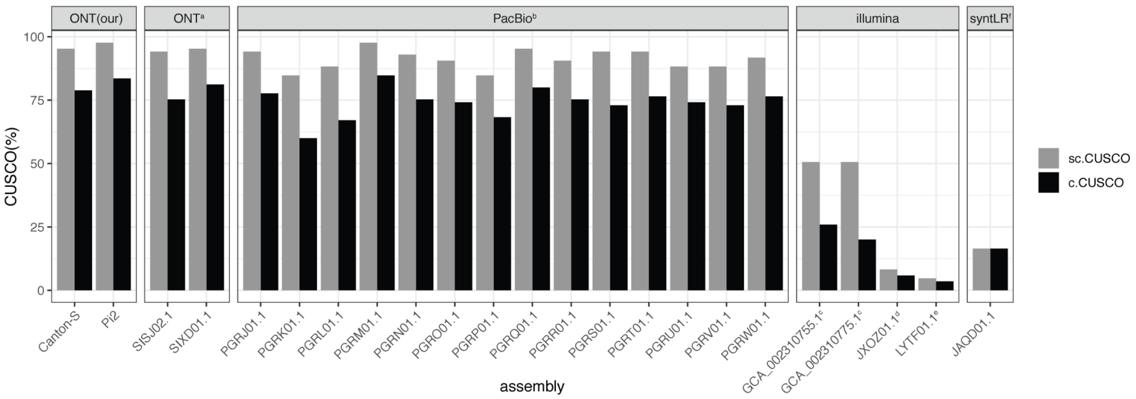


Figure 10: CUSCO values for our assemblies and publicly available assemblies of different *D. melanogaster* strains. We used assemblies from NCBI databases with following accession numbers: [a]WGS: SIXD01000000 and SISJ02000000 (Ellison and Cao, 2020) for ONT; [b]Bioproject: PRJNA418342 (Chakraborty et al., 2019) for PacBio; [c]Genbank: GCA_002310755.1 and GCA_002310775.1 (Anreiter et al., 2017), [d]WGS: JXOZ01000000 (Vicoso and Bachtrog, 2015),[e]WGS: LYTF01000000 (Singhal et al., 2017) for illumina; [f]WGS: JAQD01000000 (McCoy et al., 2014) for illumina synthetic long reads.

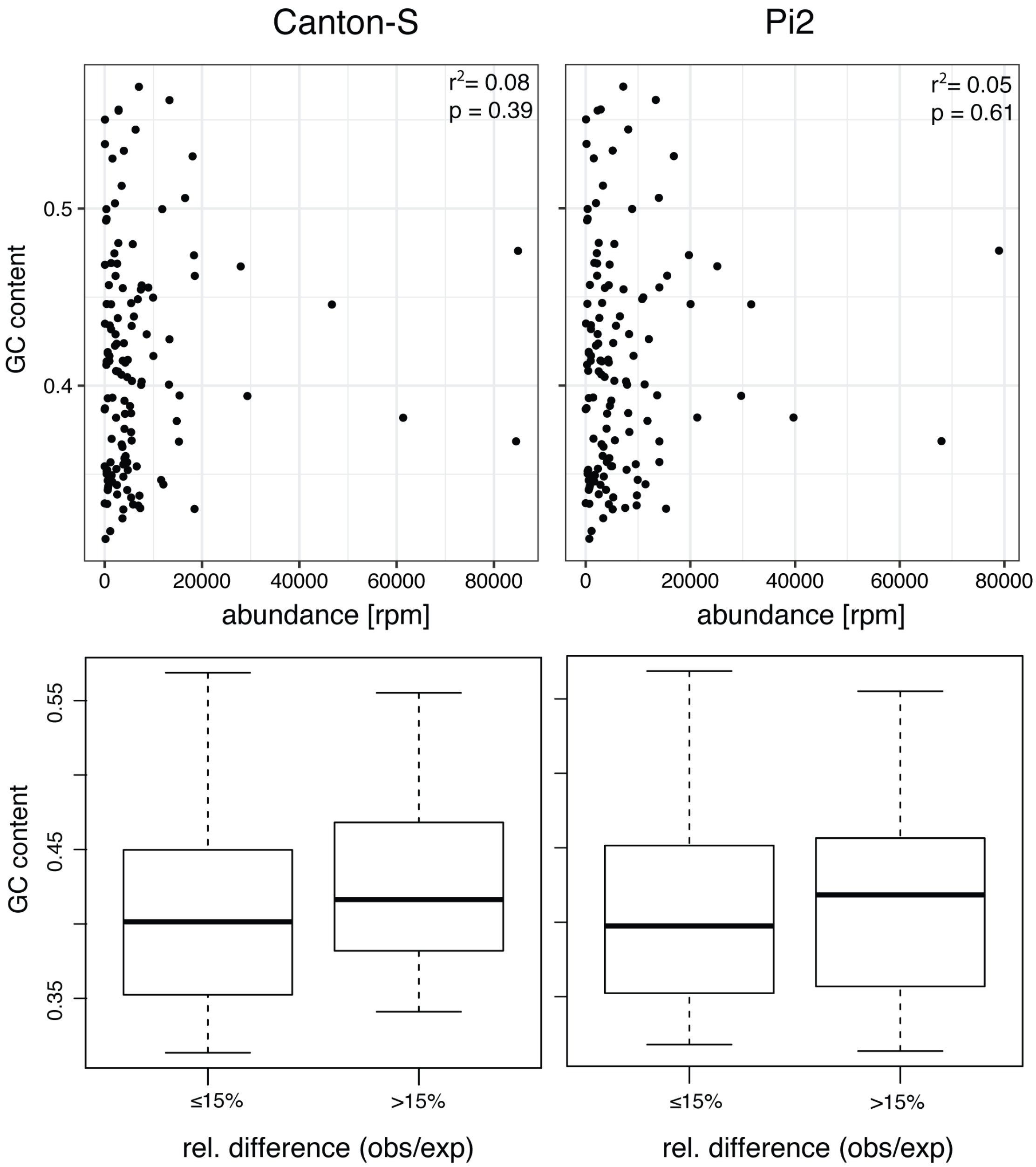


Figure 11: Influence of the GC-content on TE abundance for our assemblies of Canton-S and Pi2. A) Relationship between the the GC-content of a TE and the estimated abundance of a TE. Correlations were calculated with the spearman method. B) GC-content of TEs that strongly deviate from the expected abundance (difference between observed and expected TE abundance > 15%) compared to TEs that do not deviate from expectations (difference < 15%). For both Canton-S and Pi2 the differences between deviating and not-deviating TEs was not significant (Wilcoxon rank-sum test, two-sided: $p_{CS} = 0.109$, $p_{Pi2} = 0.3813$)

### 2 Supplementary tables

Table 1: Overview of the raw data used for the assemblies; PE paired ends

| | CantonS | Pi2 |
|---|---|---|
| ONT, coverage | 149x | 199x |
| ONT, flow cells | 2 | 3 |
| ONT, mean read length | 7146bp | 8045bp |
| Illumina PE, coverage | 30x | 40x |
| Illumina PE, read length | 125 | 125 |
| Hi-C, coverage | 591x | 260x |

Table 2: Influence of different polishing steps on the quality of Canton-S assemblies (30x coverage with long reads). Assembly quality is estimated with BUSCO and our four TE centered quality metrics (CUSCO, TE abundance, SNPs and IDs in TEs).

| quality | raw | racon | pilon |
|---|---|---|---|
| BUSCO | 76.8 | 85.6 | 98.4 |
| Cusco | 60.0 | 60.0 | 60.0 |
| TE abu. | 0.97 | 0.97 | 0.97 |
| TE SNPs | 0.93 | 0.99 | 1.00 |
| TE IDs | 0.91 | 0.97 | 0.99 |

Table 3: Effect of polishing on BUSCO values. We applied three rounds of polishing with Racon, picked the assembly with the highest BUSCO value (bold) and polished this assembly three times with Pilon, where we again kept the assembly with the highest BUSCO values (bold). In case BUSCO values did not improve between two successive iterations we kept the assembly requiring the fewest polishing steps. The finally used polishing strategy (polishing s.) for each assembly is shown at the bottom (R .. Racon, P.. Pilon). All ONT reads were used for these assemblies.

| | CantonS | | | Pi2 | | |
|---|---|---|---|---|---|---|
| | Canu | miniasm | wtdbg2 | Canu | miniasm | wtdbg2 |
| unpolished | 83.0 | 1.1 | 74.9 | 87 | 0.8 | 76.1 |
| 1x Racon | 91.0 | 80.2 | 90.5 | 93.0 | 84.2 | 92.3 |
| 2x Racon | 91.0 | 90.2 | 91.4 | 92.9 | 90.3 | **92.9** |
| 3x Racon | **91.6** | **91.5** | **91.7** | **93.3** | **93.3** | 92.6 |
| 1x Pilon | 98.6 | 98.2 | 98.5 | 98.2 | 98.3 | 96.9 |
| 2x Pilon | 98.8 | 98.6 | **98.9** | 98.4 | **98.5** | **97.7** |
| 3x Pilon | **98.9** | **98.8** | 98.8 | **98.4** | 98.5 | 97.4 |
| polishing s. | 3R,3P | 3R,3P | 3R,2P | 3R,3P | 3R,2P | 3R,2P |

Table 4: BUSCO values for assemblies generated with different assemblers and coverages for Canton-S. For each assembly the optimized number of polishing rounds performed with Racon (R) and Pilon (P) are shown (for optimization procedure see supplementary table 3).

| | coverage | 20x | 30x | 50x | 75x | 100x | 120x | 150x |
|---|---|---|---|---|---|---|---|---|
| Canu | polishing | 3R,3P | 3R,3P | 3R,3P | 2R,3P | 3R,2P | 3R,3P | 3R,3P |
| | BUSCO | 98.4 | 98.4 | 98.6 | 98.6 | 98.6 | 98.8 | 98.9 |
| miniasm | polishing | 3R,3P | 3R,2P | 3R,3P | 3R,2P | 3R,3P | 3R,3P | 3R,3P |
| | BUSCO | 98.4 | 98.4 | 98.6 | 98.6 | 98.6 | 98.8 | 98.8 |
| wtdbg2 | polishing | 1R,3P | 2R,3P | 2R,3P | 3R,3P | 3R,2P | 2R,3P | 3R,2P |
| | BUSCO | 98.8 | 98.6 | 98.7 | 98.7 | 98.7 | 98.6 | 98.9 |

Table 5: Overview of the final assemblies of Canton-S and Pi2. The assembly quality is assessed with classic quality metrics (NG50, BUSCO) as well as our TE centered quality metrics.Misassembled contigs and scaffolds were broken manually (based on dot-plots; supplementary fig. 1). c.CUSCO contig-CUSCO, sc.CUSCO scaffold-CUSCO

| | | CantonS | Pi2 |
|---|---|---|---|
| ONT | Assembler | Canu | Canu |
| | Coverage | 100x | 100x |
| | contigs | 335 | 625 |
| | NG50 | 6.8m | 4.1m |
| | length | 148m | 169m |
| | BUSCO | 82.3 | 85.8 |
| | c.Cusco | 80.00 | 77.65 |
| | TE abu. | 1.02 | 1.04 |
| | TE SNPs | 0.95 | 0.97 |
| | TE IDs | 0.93 | 0.95 |
| pol. | strategy | 2R,2P | 2R,2P |
| | contigs | 335 | 625 |
| | NG50 | 4.6m | 4.1m |
| | length | 149m | 169m |
| | BUSCO | 98.7 | 98.3 |
| | c.Cusco | 81.18 | 83.53 |
| | TE abu. | 1.01 | 1.04 |
| | TE SNPs | 0.99 | 1.01 |
| | TE IDs | 0.99 | 1.01 |
| Hi-C | scaffolds | 266 | 483 |
| | NG50 | 21.4m | 23.6m |
| | length | 149m | 169m |
| | BUSCO | 98.7 | 98.3 |
| | sc.Cusco | 84.71 | 91.76 |
| | TE abu. | 1.01 | 1.04 |
| | TE SNPs | 0.99 | 1.01 |
| | TE IDs | 0.99 | 1.01 |
| ref. scaf. | scaffolds | 15 | 16 |
| | NG50 | 28.2m | 37.2m |
| | length | 149m | 169m |
| | BUSCO | 98.6 | 98.2 |
| | sc.Cusco | 95.29 | 97.65 |
| | TE abu. | 1.01 | 1.04 |
| | TE SNPs | 0.99 | 1.01 |
| | TE IDs | 0.98 | 1.00 |

Table 6: Overview of validated presence/absence polymorphisms in piRNA clusters. For each polymorphism, we show the primers, the cluster, the TE family, the length of the expected fragment, the position and indicate whether or not the polymorphism is present in Canton-S, Pi2 or Iso-1 (y..yes, n..no). Numbers in brackets refer to lane numbers in the PCR gels (supplementary fig. 8).

| PCR-ID | strain of prescence | Canton-S | Pi2 | Iso-1 | piRNA cluster | TE family | start | end | length | primer #1 | primer #2 |
|---|---|---|---|---|---|---|---|---|---|---|---|
| cs19L | Canton-S | y(13) | n(14) | - | cl112 | doc | 14527 | 15230 | 704 | GCAGAGAGGGAGAGCAAGAA | TTTGACTGGCCTTCTTACGG |
| cs19R | Canton-S | y(16) | n(17) | - | cl112 | doc | 9735 | 10435 | 701 | TATTCAACTGGCCCTTCCTG | AATCCCCTCGCAAGAAAACT |
| cs16L | Canton-S | y(7) | n(8) | - | cl133 | mdg1 | 15190 | 15885 | 696 | TCACGGTCGCCATGTAGTTA | TCCTCGGTTGGTCCTAATTC |
| cs16R | Canton-S | y(10) | n(11) | - | cl133 | mdg1 | 7521 | 8235 | 715 | GCTGAAAGAACCCACCGATA | GCACTTGGCTGTCACAAGAG |
| cs14R | Canton-S | y(4) | n(5) | - | cl140 | doc | 58054 | 58752 | 699 | TGCGCGTAATATATGTCGATG | GTGCTCGATCACCGATTTG |
| cs14L | Canton-S | y(1) | n(2) | - | cl140 | doc | 62711 | 63311 | 601 | ACTTATGGCTCCGAGCTGTG | CCCGAAAACCGTTAAATCAG |
| cs8R | Canton-S | y(52) | n(53) | - | cl15 | 412 | 52318 | 52917 | 600 | AGCGTGATTTGGTGATAGCC | GGTCGCCATGTAATGATGAA |
| cs8L | Canton-S | y(49) | n(50) | - | cl15 | 412 | 59624 | 60219 | 596 | CCCACTCGAAGGCAAAAGTA | GAGCCCATTTATGCCAAGTT |
| cs7R | Canton-S | y(46) | n(47) | - | cl15 | F-element | 74840 | 75448 | 609 | GCTACTGCCTGATCCGATGT | AGCGTCTCTTTCGCTTTCAG |
| cs7L | Canton-S | y(43) | n(44) | - | cl15 | F-element | 79627 | 80231 | 605 | TTCAGCGAGCACAATCAAAG | TCAAAACGCAATCGGTCATA |
| cs9R | Canton-S | y(58) | y(59) | - | cl15 | springer | 27385 | 27982 | 598 | AAGCTGCTTGTGCAAGTTGA | ACTTACGCCCCTGTATGTCG |
| cs9L | Canton-S | y(55) | y(56) | - | cl15 | springer | 30676 | 31482 | 807 | GGTCAATTTTCGCACCTACC | CCTAGCAATGGCTCAGGAAA |
| cs6R | Canton-S | y(40) | n(41) | - | cl16 | 297 | 40121 | 40727 | 607 | TACTCACGCACTTTCCATCG | TCAAAACACACCACAACAAACA |
| cs6L | Canton-S | y(37) | n(38) | - | cl16 | 297 | 47115 | 47710 | 596 | GCAGCTGGGATACGTTATGG | CCAAGGCGGTCTGTATTGTT |
| cs4R | Canton-S | y(34) | n(35) | - | cl20 | roo | 119943 | 120547 | 605 | GATGGCGTCCAAGTTTGTTT | CCTTTGGTAGGGGGAAACTG |
| cs4L | Canton-S | y(31) | n(32) | - | cl20 | roo | 129025 | 129624 | 600 | CGATAAGGCGGGGACTATTT | CCTTCGGTCAAACACCTTGT |
| cs3R | Canton-S | y(28) | n(29) | - | cl26 | juan | 78770 | 79374 | 605 | CAAATAAAGCGGCGAACTGT | GGTCACCTGTTTGGGGTAGA |
| cs3L | Canton-S | y(25) | n(26) | - | cl26 | juan | 82718 | 83336 | 619 | AGCTGCATTGAAACAGTCCA | GTTTCAGCGGATCCCAAATA |
| cs1L | Canton-S | y(19) | n(20) | - | cl6 | quasimodo | 49383 | 49885 | 503 | GCCCTTCGCTTTAAGCTTTC | AGCGTTCGCTTTGCAAATTA |
| cs1R | Canton-S | y(22) | n(23) | - | cl6 | quasimodo | 56976 | 57581 | 606 | TTGTCACCAGAATTTGGGTTC | GGCCGTTTTTAACGGAAAG |
| pi22L | Pi2 | n(179) | y(180) | - | cl1 | invader1 | 18945 | 19546 | 602 | TCTTTCCGTCCATCCGTATC | CGGCAGACAACCGTTAGAGT |
| pi22R | Pi2 | n(182) | y(183) | - | cl1 | invader1 | 22817 | 23509 | 693 | CATCAGAATGCCCTTCTTCC | CTTATTGAGTCGCGAGAAACG |
| pi25L | Pi2 | n(97) | y(98) | - | cl1 | P-element | 267796 | - | 609 | GGGATTCTGCGATTTGATTC | CTTCGGTAAGCTTCGGC |
| pi25R | Pi2 | n(100) | y(101) | - | cl1 | P-element | - | 268773 | 474 | GTGGATGTCTCTTGCCG | TTGGTTTAAGCGATGCTTTG |
| pi23 | Pi2 | n(191) | y(192) | - | cl1 | rover | 30040 | 30743 | 704 | CAAATCCAACTGCATCACCA | GTTCTCAGTTCGGGTTCGAT |
| pi24 | Pi2 | n(194) | y(195) | - | cl1 | duplication | 53683 | 55687 | 2005 | GCATTAGCTCAGGCAGGAAA | TTGCTGTTGCCATTGTTGTT |
| pi17R | Pi2 | n(224) | y(225) | - | cl130 | copia | 18413 | 19115 | 703 | AGGTCTGCTCGGTGACATTC | CCACGACCTACTCACAGCAA |
| pi17L | Pi2 | n(221) | y(222) | - | cl130 | copia | 23883 | 24580 | 698 | AATGGCCACACCTTTTATGC | TTCAAACATTATCCCCTGCAC |
| pi18L | Pi2 | n(227) | y(228) | - | cl130 | idefix | 13300 | 14003 | 704 | ATCCAAGAGAAACGCAAAGC | ATATGTCGGCGAAACAGGAG |
| pi18R | Pi2 | n(230) | y(231) | - | cl130 | idefix | 5858 | 6371 | 514 | CCGGATGTCAAGAGGAAGAG | GGTGAGGCTTAAGCAAGGAA |
| pi15R | Pi2 | n(91) | y(92) | - | cl137 | 297 | 11479 | 11880 | 402 | GCAAACACACGAAATCGAAG | TCCAGCGCTCAAGTAATACG |
| pi15L | Pi2 | n(88) | y(89) | - | cl137 | 297 | 2696 | 3085 | 390 | CTGTCACGGTCGCCATATAA | TAGCGTTGAAAGAGGGCAGT |
| pi12R | Pi2 | n(79) | y(80) | - | cl140 | F-element | 28851 | 29541 | 691 | CACACAAACGGTCCATATCAA | CCGTTTATTGCTAGCGGTTT |
| pi12L | Pi2 | n(76) | y(77) | - | cl140 | F-element | 33426 | 34123 | 698 | CAACGCAATCACGGAACTTA | TTGACCCTTTGGTGGGAATA |
| pi13R | Pi2 | n(85) | y(86) | - | cl140 | F-element | 3919 | 4620 | 702 | TTCGGATTGACATTGGTTGA | TATTGACCGCTCGATGTCTG |
| pi13L | Pi2 | n(82) | y(83) | - | cl140 | F-element | 8614 | 9318 | 705 | TTCAGCGAGCACAATCAAAG | AACAAACGTGTGTGGCGTTA |
| pi11R | Pi2 | n(73) | y(74) | - | cl140 | gypsy | 69269 | 69959 | 691 | CGAAACAATCGCCATCCTAT | GGCTCATTGCCGTTAAACAT |
| pi11L | Pi2 | n(70) | y(71) | - | cl140 | gypsy | 76745 | 77449 | 705 | GGCGATAGCGATTTGATTGT | AACGCTCCACGTTTCATAGG |
| pi10R | Pi2 | n(67) | y(68) | - | cl15 | F-element | 77867 | 78569 | 703 | CTTCTCCGTGCTATCCTTCG | CTCCGCTCAGCCTAGATCAC |
| pi10L | Pi2 | n(65) | y(65) | - | cl15 | F-element | 80809 | 81505 | 697 | TCTCCAGCTGTTGTTGTTGG | TATGGATGCCTGAAGGGCTA |
| pi28L | Pi2 | n(197) | y(198) | - | cl16 | mdg1 | 21932 | 22610 | 679 | CGACTCCCACAACTTCATCA | AGCACTTGGCTGTCACAAGA |
| pi28R | Pi2 | n(200) | y(201) | - | cl16 | mdg1 | 29643 | 30335 | 693 | CTCTTGTGACAGCCAAGTGC | GAGAATCGAAACGGAAATCG |
| pi5R | Pi2 | n(188) | y(189) | - | cl19 | gypsy5 | 12115 | 12808 | 694 | AAGTAGTGGCGTGGACTGCT | TATACCGGGGAAGTGACTGG |
| pi5L | Pi2 | n(185) | y(186) | - | cl19 | gypsy5 | 4604 | 5309 | 706 | TGTTCTGACCCACTCCACTG | GGCCGTCACTTTATTCAAGG |
| pi2R | Pi2 | n(206) | y(207) | - | cl26 | mdg3 | 75530 | 76216 | 687 | GAGGCATCCGTTTGGTAAAA | AGGACGGCCGAGTTGTAGTA |
| pi2L | Pi2 | n(203) | y(204) | - | cl26 | mdg3 | 81089 | 81790 | 702 | GGTTGTCAGCGAAAAGTCGT | CACAATGTATCGCCCATCTG |
| pi21R | Pi2 | y(176) | y(177) | - | cl5 | copia | 12622 | 13018 | 397 | ATTTTCCCTTGCACGAAATG | ACCAGCACCACGACCTACTC |
| pi21L | Pi2 | n(173) | y(174) | - | cl5 | copia | 17489 | 18199 | 711 | CCAATCGAATGCTGAATGAA | AGGATCATCTGCGACTCAGG |
| pi20R | Pi2 | n(170) | y(171) | - | cl5 | gtwin | 29656 | 30354 | 699 | CACCAACGATAGCTGATCCA | GCTCGCAGCCGTAAAATATC |
| pi20L | Pi2 | n(167) | y(168) | - | cl5 | gtwin | 51179 | 51671 | 493 | CAATGCATCACGCCATAAAG | AGCAATCATGATGTCGTCCA |
| cs29R | CS | y(237) | y(236) | - | cl1 | copia | 267802 | 268194 | 393 | GTTTATTAGGCATGGACTGG | CAGCAGCAAGAATACTCC |
| cs29L | CS | y(234) | n(233) | - | cl1 | copia | 272690 | 273429 | 740 | CGCTTGTAGTCTATCCCTAAC | CCACCCACATTTGATAGTTAC |
| iso34L | Iso-1 | n(135) | n(136) | y(137) | cl1 | 1731 | 91808 | 92595 | 788 | GAATCTGTTACGGCCCATTC | CATGAAAGAGGGTCACGTTG |
| iso34R | Iso-1 | n(139) | n(140) | y(141) | cl1 | 1731 | 96320 | 97224 | 905 | TCATTCCGGCAATCTTGAGT | TGTCCGACATTGTGGGTAAT |
| iso32L | Iso-1 | n(119) | n(120) | y(121) | cl1 | F-element | 57380 | 57965 | 586 | AATGGGAATTTGTGCTTTCG | TCCTGGGCTATGGGTTATTG |
| iso32R | Iso-1 | n(123) | n(124) | y(125) | cl1 | F-element | 61836 | 62648 | 813 | CTCCGCTCAGCCTAGATCAC | CTTTGGAGGCAAATTTCCAA |
| iso35L | Iso-1/Pi2 | n(143) | y(144) | y(145) | cl1 | F-element | 131490 | 132095 | 606 | CGAACTCCATCCCATTCCTA | CCTACCAACCCAGCGAATAA |
| iso35R | Iso-1/Pi2 | y(147) | y(148) | y(149) | cl1 | F-element | 136291 | 136888 | 598 | TCCTGGGCTATGGGTTATTG | ATTTGGTCTGGTGACGCTCT |
| iso38L | Iso-1 | n(209) | n(210) | y(211) | cl1 | F-element | 239670 | 240262 | 593 | TTGTCCAGAGTGACCCTTCC | CACTGGCTCATCAACTCTGG |
| iso38R | Iso-1 | n(213) | n(214) | y(215) | cl1 | F-element | 244111 | 245067 | 957 | TCCTGGGCTATGGGTTATTG | TGTGCTCACCATAAAATGTGG |
| iso37L | Iso-1/Pi2 | n(159) | y(160) | y(161) | cl1 | invader3 | 215863 | 216462 | 600 | GCTACGCCTGTGGACAACTT | TGCTGCTGATCCGTTATTTG |
| iso37R | Iso-1/Pi2 | n(163) | y(164) | y(165) | cl1 | invader3 | 221147 | 221746 | 600 | GCCGAGTTGTGGTAGGATGA | CCCAAGGTGTACGGGTCTAA |
| iso36L | Iso-1 | n(151) | n(152) | y(153) | cl1 | juan | 156015 | 156659 | 645 | TTCACAATGCATGCCGTAAT | CCTGTTTGGGGTAGATGTGC |
| iso36R | Iso-1 | n(155) | n(156) | y(157) | cl1 | juan | 160009 | 160613 | 605 | AGCTGCATTGAAACAGTCCA | CACGACAGAGAGGGATGACA |
| iso33L | Iso-1/Pi2 | n(127) | y(128) | y(129) | cl1 | stalker4 | 75075 | 75679 | 605 | CCTCGTAGACATCGGACTGC | AAGTGGTCAATGCCTTCTGC |
| iso33R | Iso-1/Pi2 | n(131) | y(132) | y(133) | cl1 | stalker4 | 82606 | 83307 | 702 | GCAGAAGGCATTGACCACTT | GACCACCTGACGATCAAACC |
| cs30R | Canton-S | y(107) | n(108) | n(109) | cl1 | doc | 148642 | 149240 | 599 | ATACGAATGAGACCCGCAGT | CGTGGTACCTTCGAAACGA |
| cs30L | Canton-S | y(103) | n(104) | n(105) | cl1 | doc | 153030 | 153629 | 600 | CGGTCAGTGCTGCTGAATTA | ATTACGGCATGCATTGTGAA |
| cs31R | Canton-S | n(115) | y(116) | n(117) | cl1 | roo | 137094 | 137698 | 605 | ATACGAATGAGACCCGCAGT | TTGGGCTCCGTTCATATCTT |
| cs31L | Canton-S | y(111) | n(112) | n(113) | cl1 | roo | 145714 | 146422 | 709 | GGGCACATCTGCCTATCTTG | AACGGGAATGTGTGGCTATT |
